# FBXL21 regulates diurnal proteostasis and stress response by targeting DNAJB6 and client proteins

**DOI:** 10.64898/2026.05.20.726545

**Authors:** Ji Ye Lim, Jaebok Wi, Marvin Wirianto, Chorong Han, Sun Young Kim, Jane Nguyen, Sehyun Jung, Kristin Eckel-Mahan, Sung Yun Jung, Karyn A. Esser, Zheng Chen, Seung-Hee Yoo

## Abstract

Circadian regulation of proteostasis, a key determinant of muscle health, remains poorly understood. Here, we identified DNAJB6, an Hsp40 (DnaJ) co-chaperone, as a substrate of the circadian E3 ligase FBXL21. FBXL21 mediated the ubiquitination-dependent proteasomal degradation of both DNAJB6 and its client proteins including Desmin; causative mutations of DNAJB6 in myopathies, however, rendered resistance to FBXL21-directed degradation. *Fbxl21* KO C2C12 cells displayed aberrant accumulation of Desmin, and showed aggravated cytoplasmic accumulation of TDP-43, another DNAJB6 client protein, in heat shock response. Under timed exercise as a physiological stressor, WT mice displayed robust diurnal rhythms in the levels of stress granule markers (G3BP1 and FUS) and TDP-43 as a function of exercise timing. In contrast, the *Fbxl21* hypomorph *Psttm* mutant mice showed elevated expression of these proteins without exercise, which was exacerbated under exercise-induced stress conditions; importantly, these abnormalities were rescued by skeletal muscle-specific FBXL21 expression. Our study elucidates a novel diurnal regulatory mechanism of skeletal muscle proteostasis via FBXL21 as a chaperone-linked E3 ligase, highlighting the FBXL21-DNAJB6 axis as a potential therapeutic target for myopathies.

## Introduction

The circadian clock system broadly regulates daily physiology across essential organs, particularly the skeletal muscle that is mainly responsible for the daily cycle of activity^1–3^. The stark contrast between dynamic movement during the day and relative immobility during the night is accompanied by the dramatic diurnal rhythm in skeletal muscle physiology and function, underscoring a strong basis for circadian regulation^4^. At the molecular level, circadian control of muscle physiology is mediated by the ubiquitous, cell-autonomous core oscillators comprised of transcription/translation feedback loops containing positive (CLOCK, BMAL1, RORs) and negative (PERs, CRYs, REV-ERBs) core clock components^5–7^. The oscillators regulate output gene expression in a tissue-specific manner; for example, the molecular oscillators drive approximately 15-20% of the muscle transcriptome with little overlap with circadian transcriptome in other organs^8^.

Circadian gene regulation encompasses every step of the expression cascade: while transcription starts the circadian cycle, post-translational modifications and protein degradation are also essential for resetting the system for the next cycle^9–13^. Since skeletal muscle is the largest protein reservoir in our body, it is important to understand circadian regulatory mechanisms of proteostasis, the maintenance of protein homeostasis as a result of coordinated protein synthesis, chaperone-assisted folding, and degradation^14–17^. Our previous work identified the circadian E3 ubiquitin ligase FBXL21^18, 19^ as a key regulator of time-of-day-dependent protein degradation in the skeletal muscle, particularly targeting sarcomeric proteins such as TCAP and MYOZ1^20, 21^. Furthermore, recent studies also suggest a role of the clock in protein quality control via molecular chaperones. For example, several core clock components (PER2, BMAL1) have been shown to physically and functionally interact with molecular chaperones such as HSP70 and HSP90 in different cell types, suggesting a key mechanism for circadian regulation of proteostasis^22–24^.

Molecular chaperones and their regulatory cochaperones safeguard cellular proteostasis by assisting in protein folding, preventing aggregation, and coordinating with degradation pathways to preserve muscle function and integrity^24–26^. For example, HSP70 interacts with its associated E3 ligase CHIP to either facilitate the refolding of misfolded proteins or direct them toward ubiquitin-dependent proteasomal degradation. This interaction represents a crucial checkpoint in proteostasis, allowing the cell to triage damaged proteins between refolding and clearance. DNAJB6, as a member of the Hsp40 (DnaJ) family, is an essential co-chaperone of HSP70 ^27, 28^. DNAJB6 brings client proteins, including many that are aggregation-prone and involved in diseases, to HSP70 for refolding. In particular, DNAJB6 plays an important role in maintaining proteostasis under stress conditions^27, 28^. Several recent studies highlight a key role of DNAJB6 in the dynamic formation-dissolution cycles of stress granules for handling aggregated or misfolded proteins^29–32^. It directly binds to aggregation-prone substrates with amyloidogenic sequences and prevents their self-assembly into insoluble aggregates. In these cases, DNAJB6 loss of function contributes to the formation of pathological aggregates and muscle degeneration ^29–32^. Of note, several mutations of DNAJB6 in the G/F-domain were recently found to be causative mutations for Limb-Girdle Muscular Dystrophy (LGMD1D) and these mutant DNAJB6 proteins form unresolved protein aggregates in patient skeletal muscle^33^, suggesting a potential role of stress granules in this debilitating myopathy^34, 35^.

Here, we report that FBXL21 directly targets DNAJB6 as a novel substrate, promoting its ubiquitination and regulating its diurnal oscillation in skeletal muscle. FBXL21 mediates the ubiquitination and degradation of DNAJB6 client proteins, functioning as a partner E3 ligase to regulate proteostasis. Under stress conditions, DNAJB6 and FBXL21 are co-localized in heat-shock-induced stress granules for client protein degradation. Using timed exercise as a stressor model in skeletal muscle physiology, we observed a robust time-dependent exercise effect on DNAJB6 client protein accumulation in WT skeletal muscle; however, in *Fbxl21*-deficient *Psttm* mouse muscle, client protein levels were elevated under the control condition without exercise and remained consistently increased with exercise compared to WT mice. These findings suggest that the FBXL21-DNAJB6 axis regulates skeletal muscle proteostasis via diurnal degradation of client proteins and stress-induced inclusion dissolution, illustrating a novel time-of-day-dependent mechanism that modulates cellular stress response.

## Results

### Identification of DNAJB6 as a novel target of FBXL21

To investigate the role of FBXL21 in the skeletal muscle, we performed two independent unbiased screens, namely affinity pull-down using a Flag-Fbxl21 followed by mass-spectrometry (Table EV1) and yeast two-hybrid (Y2H) using a human skeletal muscle library, to identify FBXL21 targets (Fig. EV1A)^21^. Among the shared FBXL21 substrates, we focused on DNAJB6 due to its role as an HSP70 co-chaperone in muscle proteostasis and its link to limb-girdle muscular dystrophy^33, 36–41^. DNAJB6a and DNAJB6b are splice variants of the DNAJB6 gene, and the DNAJB6b form is known to play a critical role in cytoplasmic proteostasis^42^. To validate DNAJB6 as a possible FBXL21 substrate, FBXL21 and DNAJB6a or DNAJB6b were co-expressed, and co-immunoprecipitation was performed. Both DNAJB6a and DNAJB6b physically interacted with FBXL21 (Fig. 1A,B), as did the human form of DNAJB6 (Fig. EV1B,C). Interestingly, DNAJB6 displayed a diurnal pattern of expression, and co-immunoprecipitation revealed *in vivo* interaction with endogenous FBXL21 in gastrocnemius muscles collected from wild-type (WT) C57BL/6 mice at both ZT6 and ZT18 (Zeitgeber times, 6 hours after light on and light off respectively) (Fig. 1C). Next, we examined the effect of the *Fbxl21* hypomorph mutation (*Psttm*)^18, 20, 21^ on DNAJB6 diurnal rhythms in skeletal muscle. Both DNAJB6a and DNAJB6b showed robust diurnal rhythms in WT (Fig. 1D). In *Psttm* mice, DNAJB6a levels were increased at ZT6, dampening the diurnal rhythm relative to WT; in comparison, DNAJB6b levels were similarly increased at both time points compared with WT (Fig. 1D). We next performed qPCR to examine *Dnajb6* mRNA levels in WT and *Psttm* mice (Fig. EV1D). *Dnajb6* mRNA levels were comparable between WT and *Psttm* mice and also between the two time points, suggesting that the diurnal patterns of DNAJB6 protein level and its increased abundance in *Psttm* mice are primarily due to post-transcriptional regulatory mechanisms. Next, soluble and insoluble fractions were prepared from gastrocnemius muscles from WT and *Psttm* mice followed by immunoblot analysis. In the soluble fractions, both DNAJB6a and DNAJB6b levels displayed diurnal rhythms in WT and *Psttm* mice, with higher levels generally observed in *Psttm* (Fig. EV1E). In the insoluble fractions, the diurnal rhythm of DNAJB6a in WT mice was abolished in *Psttm* mice. In comparison, insoluble DNAJB6b exhibited diurnal rhythms in both WT and *Psttm* mice, with exaggerated levels detected in *Psttm* mice. Together, these findings show that FBXL21 regulates DNAJB6 turnover, and loss of FBXL21 function results in DNAJB6 accumulation.

**Fig 1.**
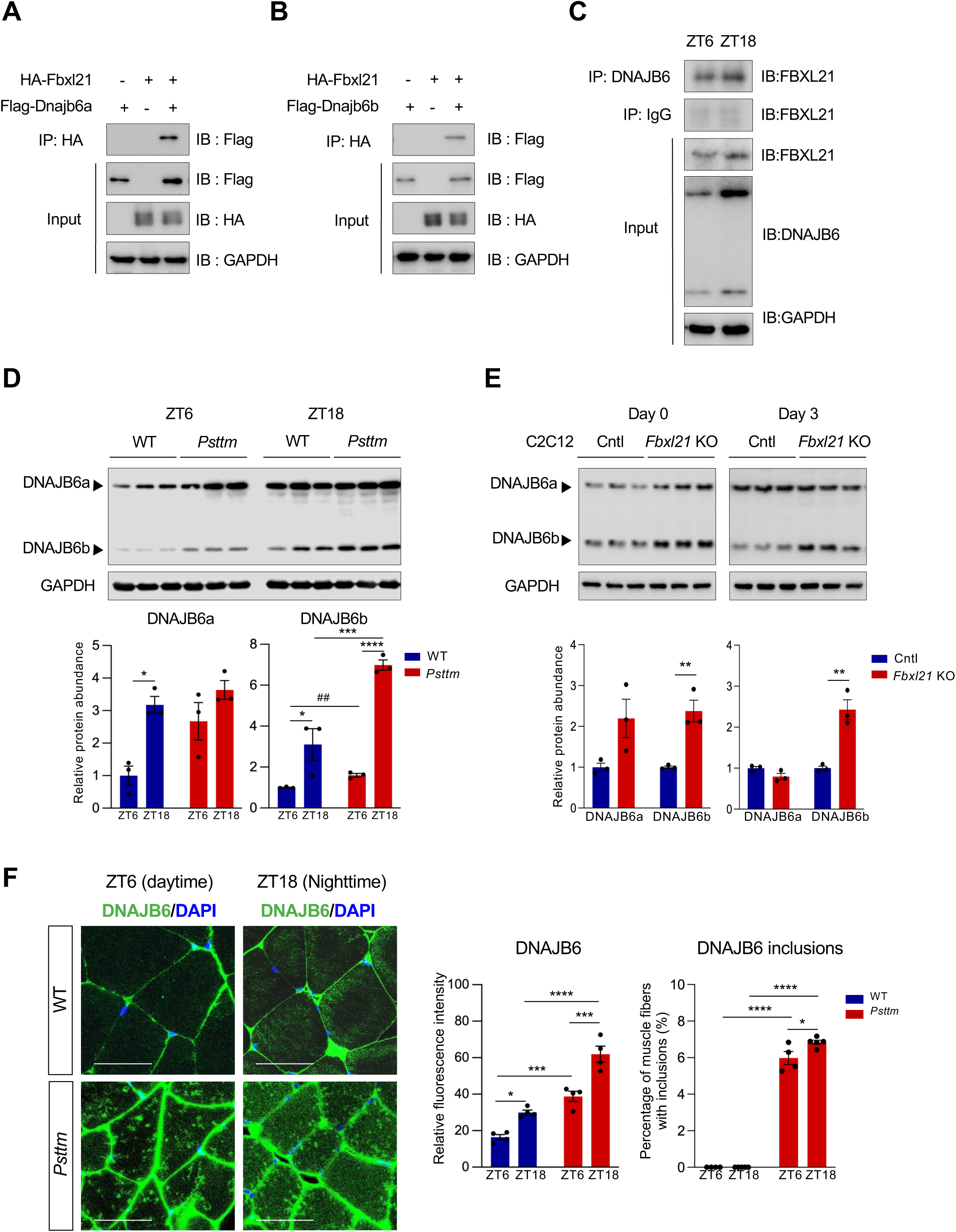
DNAJB6 is a novel target of FBXL21. Interaction of FBXL21 with DNAJB6a (**A**) and DNAJB6b (**B**). (**C**) Interaction of DNAJB6 with FBXL21. Co-immunoprecipitation from gastrocnemius tissues of WT C57BL/6J mice at the indicated Zeitgeber times (ZT), using an anti-DNAJB6 antibody. (**D**) Immunoblot analysis of DNAJB6a and DNAJB6b protein levels in gastrocnemius muscle tissues from WT and *Psttm* mice collected at ZT6 (daytime) and ZT18 (nighttime). Data are presented as mean ± SEM (n = 3 mice/group). No mark indicates not statistically significant. \**P* = 0.0145 (DNAJB6a), \**P* = 0.0270 (DNAJB6b), \*\*\**P* = 0.0007*, ****P* < 0.0001; Two-way ANOVA with Tukey’s multiple comparisons indicating a significant difference between WT and *Psttm* mice or between two time points (ZT6 and ZT18). ^##^*P* = 0.024; Unpaired Student’s *t*-test indicating a significant difference between WT and *Psttm* mice. (**E**) Immunoblotting analysis of DNAJB6a and DNAJB6b protein expression in control (Cntl) and *Fbxl21* KO C2C12 cells on differentiation Day 0 and Day 3. Data are presented as mean ± SEM (n = 3). \*\**P* = 0.0070 (Day 0), \*\**P* = 0.0045 (Day 3); Unpaired Student’s *t*-test indicating a significant difference between Cntl and *Fbxl21* KO C2C12 cells. (**F**) Immunofluorescence staining of DNAJB6 in gastrocnemius muscle cross-sections from WT and *Psttm* mice at ZT6 and ZT18. Right panels: DNAJB6 fluorescence intensity quantification and DNAJB6 inclusions in more than 1,500 muscle fibers per mouse. Data are presented as mean ± SEM (n = 4–5 mice/group), \**P* = 0.0211 (DNAJB6 fluorescence intensity), \**P* = 0.0151 (DNAJB6 inclusions), \*\*\**P* = 0.0003 (*Psttm* ZT6 vs. *Psttm* ZT18), \*\*\**P* = 0.0005 (WT ZT6 vs. *Psttm* ZT6), \*\*\*\**P* < 0.0001; Two-way ANOVA with Tukey’s multiple comparisons indicating a significant difference between WT and *Psttm* mice or between two time points (ZT6 and ZT18) in *Psttm* mice. Scale bar: 50 μm. N numbers indicate biological replicates for each sample group, and representative data from 3 independent experiments are shown.

We next employed CRISPR-Cas9-mediated *Fbxl21* knockout (KO) in C2C12 cells as an *in vitro* model (Fig. EV1F,G). In undifferentiated C2C12 cells (day 0), both isoforms were significantly elevated in *Fbxl21* KO cells; following differentiation (day 3), DNAJB6b showed marked increases compared to WT (Fig. 1E). To examine the localization of DNAJB6 in skeletal muscle tissue, we performed immunofluorescent staining using WT and *Psttm* gastrocnemius muscles collected at ZT6 and ZT18 (Fig. 1F). Consistent with immunoblotting results in Fig. 1C, immunofluorescent staining of cross-sectioned gastrocnemius muscle showed that DNAJB6 amounts were significantly increased in *Psttm* mice compared to WT at both time points (Figs. 1F, EV1H). Interestingly, DNAJB6 was exclusively localized in sarcoplasmic inclusion bodies in *Psttm* mice (Fig. 1F), indicating that loss of FBXL21 leads to accumulation of damaged or dysfunctional DNAJB6, compromising protein quality control.

### FBXL21 regulates ubiquitination-dependent DNAJB6 degradation

We next performed cycloheximide (CHX) chase assays to measure DNAJB6 half-life using control and *Fbxl21* KO C2C12 cells. In *Fbxl21* KO C2C12 cells, the degradation rates of both DNAJB6 isoforms were reduced compared to those in control cells (Fig. 2A). To validate this finding, we performed a gain-of-function experiment by expressing FBXL21 in 293T cells. Based on the strong effect of FBXL21 depletion on DNAJB6b in skeletal muscle and C2C12 cells (Fig. 1D,E), we focused on the DNAJB6b isoform. Consistent with C2C12 cells (Fig. 2A,B), the expression of FBXL21 in 293T cells accelerated DNAJB6b degradation (half-life: 2.58 h vs. 5.56 h in control cells) (Fig. 2C). Treatment with MG132, a 26S proteasome inhibitor, blocked FBXL21-mediated degradation of mouse DNAJB6b, suggesting that FBXL21 may promote the ubiquitination and proteasomal turnover of DNAJB6b (Fig. 2C). FBXL21 expression also increased the rate of human DNAJB6 degradation (Fig. EV2A,B). Interestingly, the GSK-3β inhibitor CHIR-99021 decelerated FBXL21-mediated degradation, suggesting that GSK-3β may regulate FBXL21-dependent DNAJB6b degradation similar to previously reported FBXL21 target substrates^20, 21^ (Fig. 2C). In ubiquitination assays, both mouse DNAJB6 isoforms (Fig. 2D,E) and human DNAJB6 (Fig. EV2C,D) were efficiently ubiquitinated in the presence of FBXL21. To determine whether the FBXL21-mediated ubiquitination of DNAJB6b requires SCF complex assembly, we performed ubiquitination assays using an F-box deletion mutant of FBXL21 (ΔFbxl21). Deletion of the FBXL21 F-box abolished DNAJB6b ubiquitination (Fig. EV2E), indicating that SKP1 interaction and SCF complex assembly are required. Together, these results show that FBXL21 can promote the ubiquitination of DNAJB6 and its targeting to proteasomal degradation.

**Fig 2.**
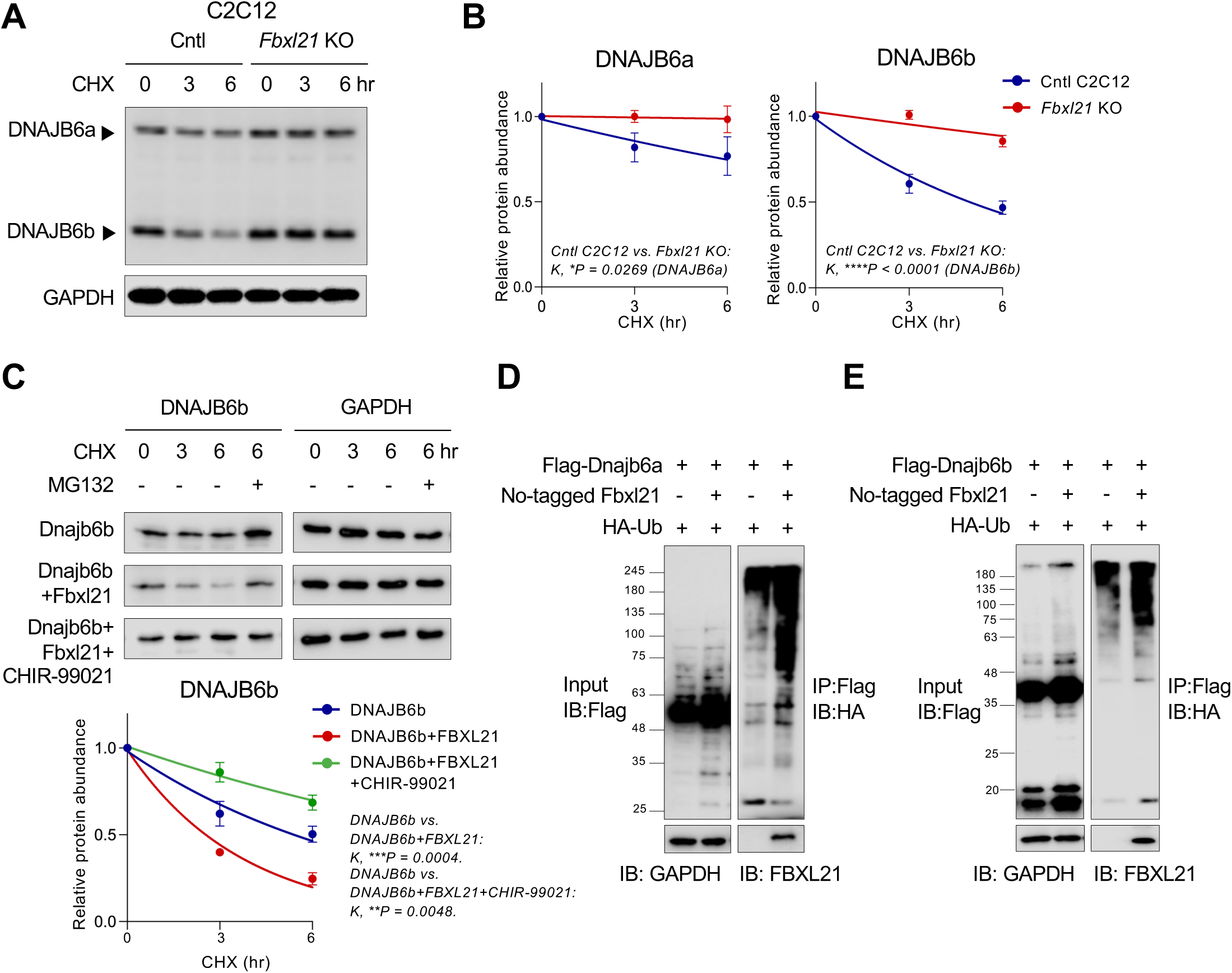
FBXL21 mediates DNAJB6 proteasomal degradation by ubiquitination. (**A-B**) Cycloheximide (CHX) chase assays assessing the stability of DNAJB6a and DNAJB6b in Cntl and *Fbxl21* KO C2C12 cells. Right panels: quantification of DNAJB6a and DNAJB6b protein stability. The half-life was calculated using nonlinear one-phase decay analysis in GraphPad Prism. In Cntl C2C12 cells, DNAJB6a and DNAJB6b half-lives are 15.12 h and 5.06 h, respectively; in *Fbxl21* KO cells, both proteins have half-lives >24.0 h (K, the half-life parameter, was significantly different in *Fbxl21* KO cells: DNAJB6a, \**P* = 0.0269; DNAJB6b, \*\*\*\**P* < 0.0001). Data are presented as mean ± SEM (n = 3–6). (**C**) The GSK-3 inhibitor CHIR-99021 blocks FBXL21-mediated degradation of DNAJB6b. 293T cells were co-transfected with the indicated constructs. Half-lives: DNAJB6b, 5.56 h; DNAJB6b/Fbxl21, 2.58 h; DNAJB6b/Fbxl21/CHIR-99021, 11.31 h. (K, significantly different between DNAJB6b and DNAJB6b/Fbxl21, \*\*\**P* = 0.0004; and between DNAJB6b and DNAJB6b/Fbxl21/CHIR-99021, \*\**P* = 0.0048). Data are presented as mean ± SEM (n = 3). (**D-E**) FBXL21 promotes ubiquitination of DNAJB6a and DNAJB6b. Flag-tagged Dnajb6a (**D**) or Flag-Dnajb6b (**E**) was co-transfected with untagged Fbxl21 and HA-tagged ubiquitin (HA-Ub), followed by ubiquitination assays. N numbers indicate biological replicates for each sample group, and representative data from 3 independent experiments are shown.

### FBXL21 requires HSP70 for stability but degrades DNAJB6 independent of CHIP

DNAJB6 is a known binding partner of HSP70^28, 43–45^; therefore, we investigated whether FBXL21 also interacts with HSP70 and mediates its degradation. Co-immunoprecipitation analysis revealed that FBXL21 interacted with HSP70 (Fig. 3A) and DNAJB6 knockdown did not affect FBXL21-HSP79 interaction, suggesting that FBXL21 can interact with HSP70 independently of DNAJB6 (Fig. 3B). However, HSP70 protein levels remained unchanged in *Psttm* mouse skeletal muscle (Fig. EV3A) and *Fbxl21* KO C2C12 cells (Fig. EV3B), suggesting that HSP70 is not a degradation target of FBXL21. Surprisingly, although HSP70 turnover was not affected by FBXL21 expression, the half-life of FBXL21 was significantly increased in the presence of HSP70 (Fig. 3C,D). This result suggests that HSP70 may stabilize FBXL21, likely functioning as its molecular chaperone. In accordance, HSP70 knock-down by siRNA strongly inhibited FBXL21-mediated DNAJB6 ubiquitination, indicating that HSP70 plays an important role in FBXL21 folding and E3 ligase activity (Fig. 3E). To determine whether DNAJB6 ubiquitination-mediated degradation is specifically regulated by FBXL21, rather than by the HSP70-associated E3 ligase CHIP, we first knocked down CHIP using siRNA and performed CHX-chase assays in 293T cells. CHIP knockdown did not affect the rate of DNAJB6 degradation (half-lives: DNAJB6b, 4.15 h; DNAJB6b/Fbxl21, 1.58 h; DNAJB6b + CHIP siRNA, 4.49 h; DNAJB6b/Fbxl21 + CHIP siRNA, 1.86 h) by FBXL21 (Fig. 3F), suggesting that CHIP may not be required for its degradation. We next treated control and *Fbxl21* KO C2C12 cells with JG-231^46, 47^, an allosteric inhibitor of HSP70 that disrupts its interaction with co-chaperones, including DNAJB6. JG-231 treatment slightly accelerated DNAJB6b degradation in control cells, whereas no significant effect on DNAJB6 degradation was observed in *Fbxl21* KO cells between control and JG-231 treatment (Fig. 3G). These results demonstrate that DNAJB6b degradation may be mediated by FBXL21, independently of CHIP.

**Fig 3.**
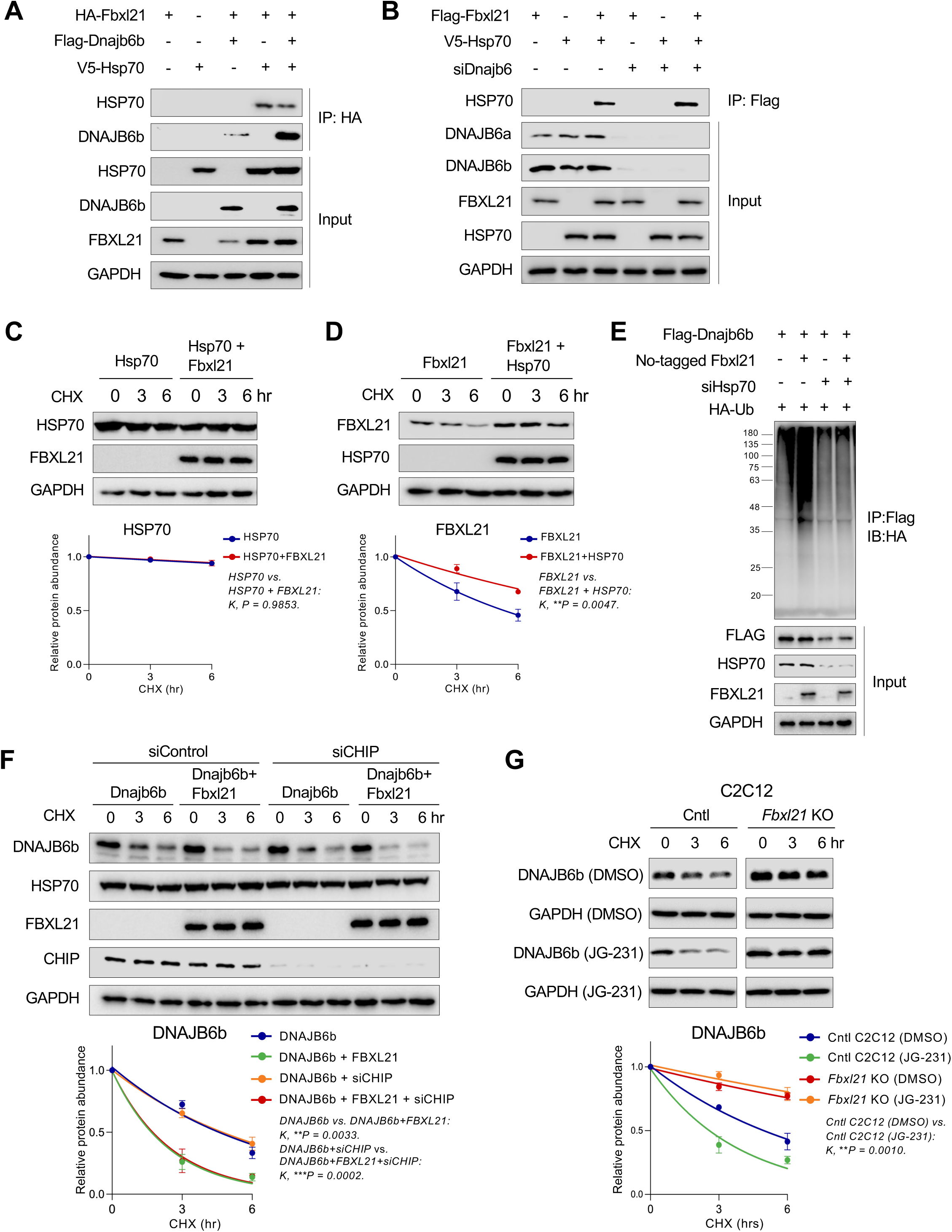
FBXL21 depends on HSP70 for its stability and facilitates DNAJB6 degradation independently of CHIP. (**A**) FBXL21 interaction with DNAJB6 and HSP70. HEK293T cells were co-transfected with the indicated constructs. Protein complexes were immunoprecipitated using anti-HA beads. (**B**) FBXL21 interacts with HSP70 independently of DNAJB6. Flag-tagged Fbxl21 and V5-tagged Hsp70 were co-transfected into 293T cells with or without Dnajb6 siRNA, followed by co-immunoprecipitation assay. (**C**) CHX chase assays assessing HSP70 protein stability. 293T cells were transfected with V5-tagged Hsp70 with or without Fbxl21. Lower panel: quantification of HSP70. HSP70 Half-lives: Hsp70, >24 h; Hsp70/Fbxl21, >24 h. (K, *P* = 0.9853). Data are presented as mean ± SEM (n = 3). (**D**) CHX chase assays assessing FBXL21 protein stability. 293T cells were transfected with Flag-tagged Fbxl21 with or without V5-tagged HSP70. Lower panel: quantification of FBXL21. FBXL21 Half-lives: Fbxl21, 5.33 h; Fbxl21/Hsp70, 11.21 h (K, significantly different between groups, \*\**P* = 0.0047). Data are presented as mean ± SEM (n = 3). (**E**) FBXL21-mediated DNAJB6b ubiquitination is inhibited by HSP70 knockdown. 293T cells were co-transfected with the indicated constructs, with or without siRNA targeting HSP70, followed by ubiquitination assays. (**F**) CHX chase assays assessing DNAJB6b stability. 293T cells were transfected with Flag-tagged DNAJB6b and/or Fbxl21, with or without CHIP knockdown (CHIP siRNA). Half-lives: DNAJB6b, 4.42 h; DNAJB6b/Fbxl21, 1.69 h; DNAJB6b + CHIP siRNA, 4.70 h; DNAJB6b/Fbxl21 + CHIP siRNA, 1.75 h (K, significantly different between DNAJB6b and DNAJB6b/Fbxl21, \*\**P* = 0.0033; and between DNAJB6b + siCHIP and DNAJB6b/Fbxl21 + siCHIP, \*\*\**P* = 0.0002). Data are presented as mean ± SEM (n = 3). (**G**) CHX chase assays assessing DNAJB6b protein stability in control and *Fbxl21* KO C2C12 treated with JG231 (an HSP70 inhibitor). Half-lives: Cntl C2C12 (DMSO), 4.91 h; Cntl C2C12 (JG-231), 2.64 h; *Fbxl21* KO C2C12 (DMSO), 15.55 h; *Fbxl21* KO C2C12 (JG-231), >24.0 h (K, significantly different between DMSO-and JG-231-treated Cntl C2C12 cells: \*\**P* = 0.0010). Data are presented as mean ± SEM (n = 3). N numbers indicate biological replicates for each sample group, and representative data from 3 independent experiments are shown.

### LGMD1D-linked DNAJB6 mutants are resistant to FBXL21-mediated degradation

Causative DNAJB6 mutations have been identified in LGMD1D patients where the mutant proteins form pathological protein inclusions in skeletal muscle. To investigate whether LGMD1D-associated DNAJB6 mutants are resistant to FBXL21-dependent proteasomal degradation, we performed co-immunoprecipitation and CHX chase assays. First, two human DNAJB6 mutations, F89I and P96R^33, 35, 36^, were individually introduced to mouse DNAJB6 as F90I and P97R (Fig. 4A). Despite similar FBXL21 binding as WT DNAJB6 (Fig. 4A), both mutants showed significantly longer half-life relative to WT in CHX chase assays (Half-lives: WT Dnajb6b, 4.83 h; WT Dnajb6b/Fbxl21, 2.35 h; Dnajb6bF90I, 11.40 h; Dnajb6bF90I/Fbxl21, 8.01 h; Dnajb6bF97R, 18.94 h; Dnajb6bF97R, 11.58 h), which was not affected by ectopic expression of FBXL21 (Fig. 4B). Moreover, both mutants showed markedly reduced ubiquitination by FBXL21 (Fig. 4C). Therefore, while the F90I and P97R mutants retain the ability to bind FBXL21 notwithstanding their structural abnormalities^38, 48, 49^, they appear resistant to FBXL21-mediated ubiquitination and proteasomal degradation, consequently prone to pathogenic accumulation. Previously, DNAJB6 mutations were shown to cause aberrant high-affinity binding to HSP70, resulting in its sequestration^47, 49^. In WT skeletal muscle, *in vivo* co-immunoprecipitation analysis revealed a diurnal pattern of DNAJB6–HSP70 binding, with stronger interaction at ZT18 than at ZT6 (Fig. 4D). In contrast, this interaction was considerably weaker in *Psttm* mice (Fig. 4D), indicating that loss of FBXL21 may impair the DNAJB6–HSP70 interaction despite the accumulation of DNAJB6.

**Fig 4.**
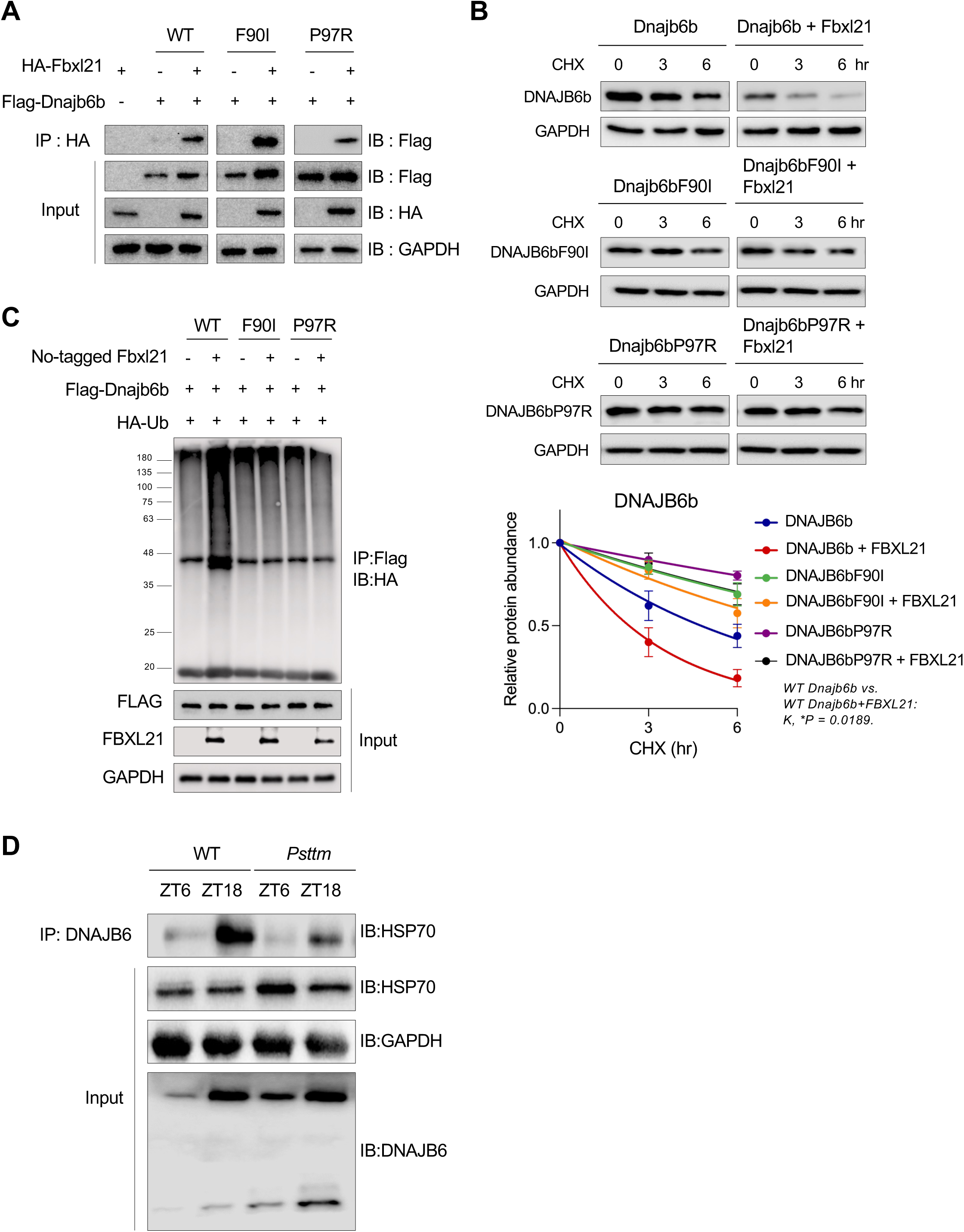
DNAJB6 mutants associated with LGMD1D exhibit resistance to FBXL21-dependent degradation. (**A**) Interaction between FBXL21, wild-type (WT) DNAJB6b, and DNAJB6 mutants. HEK293T cells were co-transfected with HA-tagged Fbxl21 and the indicated DNAJB6 constructs. Protein complexes were immunoprecipitated using anti-HA beads. (**B**) CHX chase assays assessing the stability of WT DNAJB6b and DNAJB6b mutants. 293T cells were co-transfected with Flag-tagged Fbxl21 and indicated constructs. Half-lives: WT Dnajb6b, 4.83 h; WT Dnajb6b/Fbxl21, 2.35 h; Dnajb6bF90I, 11.40 h; Dnajb6bF90I/Fbxl21, 8.01 h; Dnajb6bF97R, 18.94 h; Dnajb6bF97R, 11.58 h (K, significantly different between WT Dnajb6b and WT Dnajb6b/Fbxl21, \**P* = 0.0189; but not significant between Dnajb6b mutants and their respective Fbxl21 co-transfections). Data are presented as mean ± SEM (n = 3). (**C**) Ubiquitination assays of DNAJB6 mutants. Flag-tagged Dnajb6 and Dnajb6 mutants, untagged Fbxl21, and HA-tagged ubiquitin (HA-Ub) were co-transfected into 293T cells, followed by ubiquitination assays. (**D**) Co-immunoprecipitation of DNAJB6/HSP70 from gastrocnemius tissues of WT and *Psttm* mice at ZT6 and ZT18, using an anti-DNAJB6 antibody. N numbers indicate biological replicates for each sample group, and representative data from 3 independent experiments are shown.

### FBXL21 mediates DNAJB6 client protein degradation

To investigate whether FBXL21 plays a broader role in proteostasis by acting as a chaperone-associated E3 ligase, we next examined the degradation of DNAJB6-associated client proteins. First, we found that Desmin, a DNAJB6 client protein, strongly accumulated in *Psttm* mice skeletal muscle at both circadian time points compared to WT (Figs. 5A, EV3C). Importantly, Desmin inclusion bodies, a hallmark of Desmin-related myopathies (DRM)^50, 51^, were only observed in *Psttm* mice (Fig. 5A), suggesting a functional link between FBXL21 activity and Desmin turnover. Consistent with this observation, Desmin accumulation in both soluble and insoluble fractions was significantly elevated in *Psttm* skeletal muscle at both time points (Fig. EV3D). We observed elevated levels of Desmin and DNAJB6 in *Fbxl21* KO compared to control C2C12 cells (Figs. 5B and EV3E). Heat shock induced an increase in DNAJB6 expression in both control and *Fbxl21* KO C2C12 cells, with a more pronounced elevation in the *Fbxl21* KO C2C12 cells (Fig. 5B). Desmin levels remained higher in *Fbxl21* KO C2C12 cells under both normal and heat shock conditions (Fig. 5B), consistent with impaired degradation. To further investigate the role of FBXL21 in DNAJB6 client protein degradation, we performed co-IP and found that FBXL21 is a constituent of the *in vivo* complex of DNAJB6-HSP70-Desmin at both time points (Fig. 5C). Next, we performed CHX chase assays which revealed that loss of *Fbxl21* in C2C12 cells significantly decelerated Desmin degradation (Fig. 5D). Whereas ectopic expression of FBXL21 alone in 293T cells did not impact Desmin half-life (Desmin, 13.71 h; Desmin/Fbxl21, 13.06 h) (Fig. 5E), co-expression of DNAJB6 with FBXL21 increased Desmin degradation rate (Desmin/Fbxl21/Dnajb6b, 3.18 h) (Fig. 5E), indicating that FBXL21 may serve as an DNAJB6-linked E3 ligase to promote the degradation of DNAJB6 client proteins. Conversely, in the presence of Desmin, the degradation rate of DNAJB6 by FBXL21 was reduced (Fig. 5E), suggesting a substrate pivot when FBXL21 engages with client proteins. Finally, ubiquitination assays revealed that FBXL21-mediated Desmin ubiquitination was strongly enhanced in the presence of DNAJB6, a process dependent on HSP70 but not CHIP (Fig. 5F,G). These results indicate that FBXL21 can target both DNAJB6 and its client proteins such as Desmin, suggesting DNAJB6-FBXL21 as a novel chaperone-E3 ligase axis involved in proteostasis regulation.

**Fig 5.**
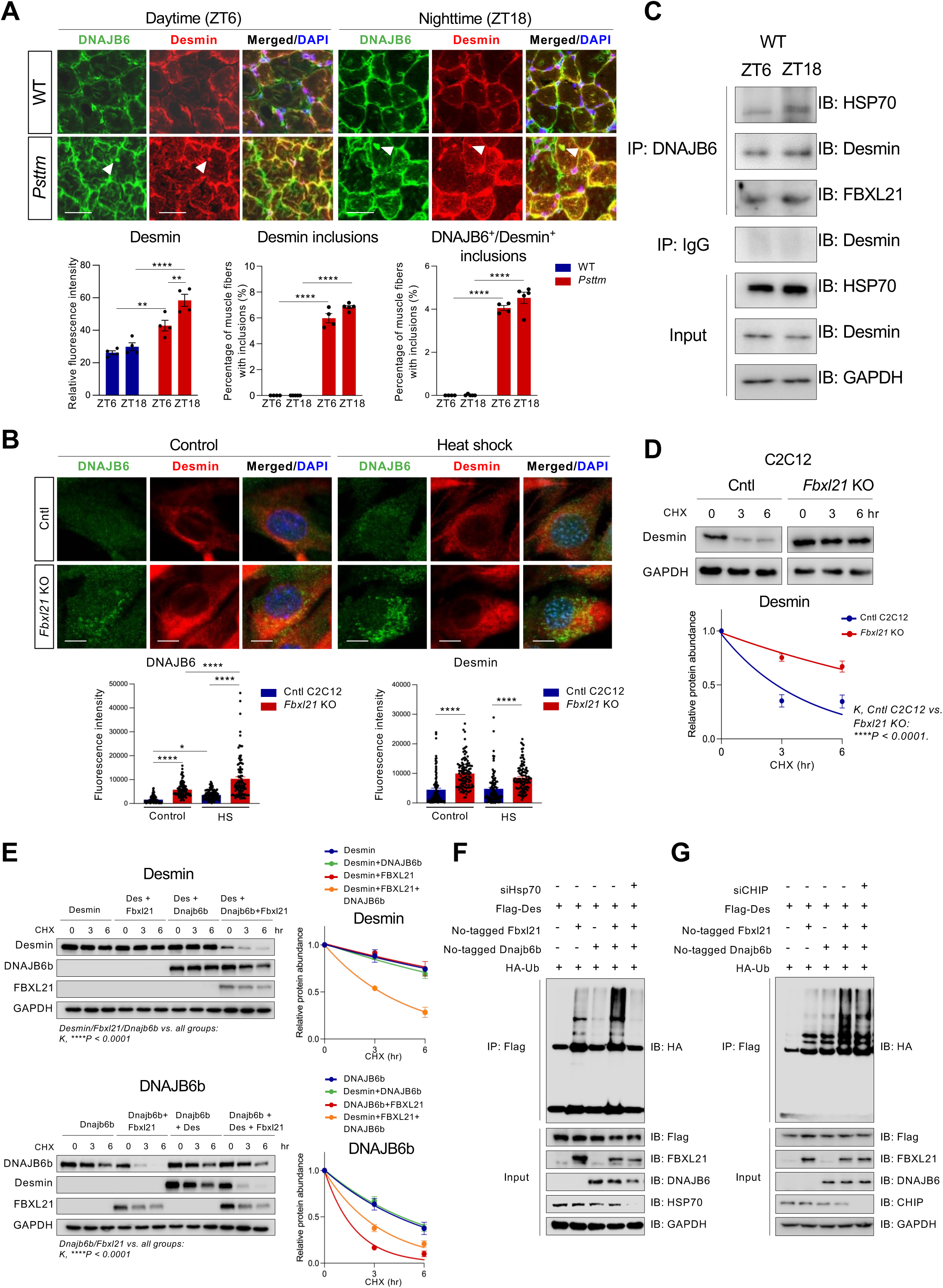
FBXL21 regulates the ubiquitination-mediated degradation of Desmin, a client protein of DNAJB6. (**A**) Immunofluorescence staining of DNAJB6 and Desmin in gastrocnemius cryosections from WT an *Psttm* mice collected at ZT6 and ZT18. Imaging was performed using confocal microscopy. Lower panels: Quantification of relative fluorescence intensity (left), percentage of muscle fibers with Desmin inclusions (middle), and percentage of muscle fibers with co-localized DNAJB6⁺/Desmin⁺ inclusions (right) (n = 4–5 mice/group/time point). No mark indicates not statistically significant. \*\**P* = 0.0064 (WT ZT6 vs. *Psttm* ZT6)*, **P* = 0.0098 (*Psttm* ZT6 vs. *Psttm* ZT18)*, ****P* < 0.0001; two-way ANOVA indicates statistical differences between genotypes (WT vs. *Psttm*) or time points (ZT6 vs. ZT18). (**B**) STED microscopy of Desmin and DNAJB6 in Cntl and *Fbxl21* KO C2C12 cells under control (37°C) and heat shock (HS; 42°C for 2 h) conditions. Lower panel: Quantification of Desmin and DNAJB6 signal intensity in Cntl and *Fbxl21* KO C2C12 cells. Data are presented as mean ± SEM (n≥100 cells/group). \**P* = 0.0136 and \*\*\*\**P* < 0.0001; One-way ANOVA with Tukey post-hoc test indicates statistical differences between groups. (**C**) Interaction of DNAJB6, HSP70, Desmin, and FBXL21 assessed by co-immunoprecipitation using gastrocnemius muscle of WT C57BL/6J mice collected at ZT6 and ZT18 with anti-DNAJB6 antibody. (**D**) CHX chase assays assessing Desmin protein stability in Cntl and *Fbxl21* KO C2C12 cells. Desmin half-lives: 2.85 h (Cntl C2C12) vs. 9.85 h (*Fbxl21* KO C2C12) (K, significantly different in *Fbxl21* KO cells, \*\*\*\**P* < 0.0001). Data are presented as mean ± SEM (n = 3). (**E**) CHX chase assays assessing Desmin and DNAJB6b stability in 293T cells co-transfected with Flag-tagged Desmin, Fbxl21, and Dnajb6b. Left panel (Desmin half-lives): Desmin, 14.20 h; Desmin/Fbxl21, 14.75 h; Desmin/Dnajb6b, 11.63 h; Desmin/Fbxl21/Dnajb6b, 3.33 h (K, Desmin/Fbxl21/Dnajb6b significantly different from all other groups, \*\*\*\**P* < 0.0001). Right panel (Dnajb6b half-lives): Dnajb6b, 4.35 h; Dnajb6b/Fbxl21, 1.26 h; Dnajb6b/Desmin, 4.57 h; Dnajb6b/Desmin/Fbxl21, 2.35 h (K, Dnajb6b/Fbxl21 significantly different from all other groups, \*\*\*\**P* < 0.0001). Data are presented as mean ± SEM (n = 4). (**F**) DNAJB6b enhances FBXL21-mediated ubiquitination of Desmin, which is mitigated by HSP70 knockdown. Flag-tagged Desmin, untagged Fbxl21, untagged Dnajb6b, and HA-tagged ubiquitin (HA-Ub) were co-transfected into 293T cells with or without Hsp70 siRNA, followed by ubiquitination assays. (**G**) CHIP knockdown does not affect the FBXL21/DNAJB6b-mediated ubiquitination of Desmin. Flag-tagged Desmin, untagged Fbxl21, untagged Dnajb6b, and HA-Ub were co-transfected with or without CHIP siRNA into 293T cells, followed by ubiquitination assays. N numbers indicate biological replicates for each sample group, and representative data from 3 independent experiments are shown.

DNAJB6 is well known to suppress polyQ aggregation by acting as a molecular chaperone to maintain the solubility of polyglutamine-expanded proteins^34, 52, 53^. We next examined whether FBXL21 depletion affects DNAJB6 activity to suppress polyQ aggregation. We introduced expression of GFP-polyQ74, as a canonical expanded polyQ, to control and *Fbxl21* KO C2C12 cells and monitored GFP inclusions after heat shock. While largely dissolved 2 h after heat shock in control C2C12 cells, abundant polyQ inclusions remained undissolved in *Fbxl21* KO cells (Fig. EV3F), suggesting a requisite role of FBXL21 in DNAJB6 chaperone function. As a negative control, we used the non-expanded polyQ construct GFP-polyQ23 and found no aggregation in both control and *Fbxl21* KO C2C12 cells, confirming that the observed aggregation was specific to polyQ-expanded proteins (Fig. EV3F).

### Fbxl21 deficiency impairs stress granule resolution and aggravates aberrant TDP-43 accumulation in C2C12 cells

In response to stress, DNAJB6 localizes to stress granules, the cytoplasmic foci containing untranslated mRNAs and misfolded proteins, where it acts with HSP70 to promote their disassembly; disease mutations in DNAJB6 impair this process^54^. To determine whether FBXL21 also localizes to stress granules under stress conditions, we performed immunofluorescence staining for endogenous FBXL21 in C2C12 cells. Upon heat shock, FBXL21 co-localized with both DNAJB6 in cytoplasmic puncta (Fig. 6A) and the stress granule marker G3BP1 (Fig. 6B) in control C2C12 cells. Due to overlapping host species of the available antibodies detecting endogenous proteins, these co-localization experiments were performed pair-wise rather than all three simultaneously. Interestingly, in *Fbxl21* KO C2C12 cells, G3BP1-positive stress granules were detected under the control condition, and G3BP1 levels were markedly elevated after heat shock compared to control cells (Fig. 6C). Consistent with the endogenous protein co-localization observed in control C2C12 cells, ectopic expression of Flag-tagged Dnajb6b and Fbxl21, along with G3BP1-RFP, revealed that DNAJB6b and Fbxl21 colocalized in response to heat shock (Fig. EV4). These observations suggest that FBXL21 regulates the formation of stress-induced protein inclusions.

**Fig 6.**
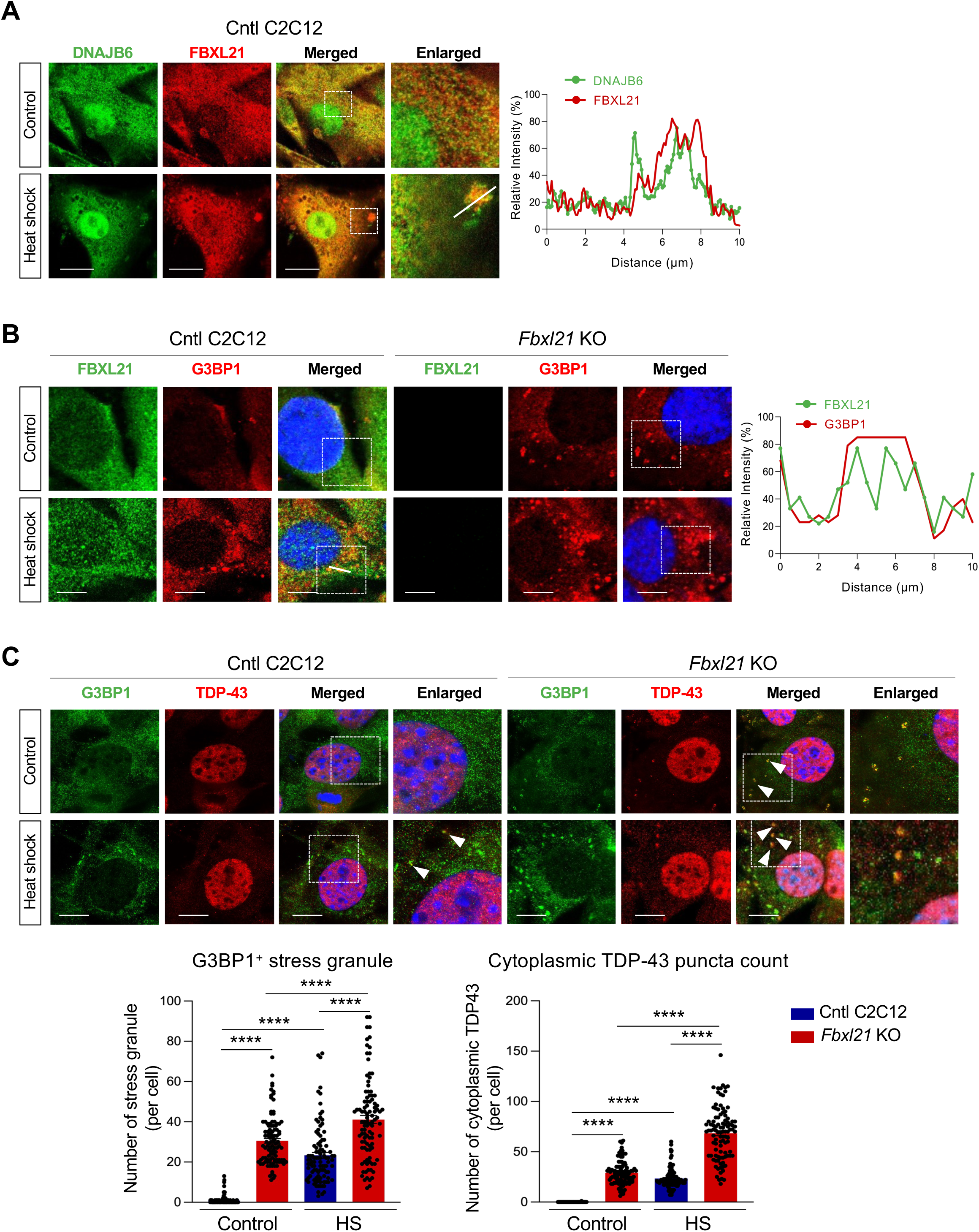
FBXL21 regulates stress granule formation and cytoplasmic TDP-43 accumulation. (**A**) Immunofluorescence staining of endogenous DNAJB6 and FBXL21 in Cntl C2C12 cells, visualized by confocal microscopy. Right panel: Line profiles showing the mean fluorescence intensities of DNAJB6 (green) and FBXL21 (red) under heat shock conditions in Cntl C2C12 cells, analyzed using Prism. Scale bar: 20 μm. (**B**) Immunofluorescence staining of FBXL21 and G3BP1 in Cntl and *Fbxl21* KO C2C12 cells, visualized by confocal microscopy. Right panel: Line profiles showing the mean fluorescence intensities of FBXL21 (green) and G3BP1 (red) under heat shock conditions in Cntl C2C12 cells, analyzed using Prism. (**C**) STED microscopy analysis of TDP-43 and G3BP1 (a stress granule marker) in Cntl and *Fbxl21* KO C2C12 cells. Lower left panel: Quantification of G3BP1^+^ stress granules per cell. Lower right panel: Cytoplasmic TDP-43 per cell. Data are presented as mean ± SEM (n≥100 cells/group). No mark indicates not statistically significant. \*\*\*\**P* <0.0001; One-way ANOVA showing significant differences between Cntl and *Fbxl21* KO C2C12 cells or between control and stressed (HS) conditions. Scale bar: 10 μm. N numbers indicate biological replicates for each sample group, and representative data from 3 independent experiments are shown.

Stress granules transiently sequester aggregation-prone proteins, including RNA-binding proteins such as TDP-43, also a client of DNAJB6; TDP-43 cytoplasmic accumulation and aggregation is a well-documented pathological hallmark of many diseases, including myopathy^49, 55–57^. To examine whether FBXL21 regulates TDP-43 inclusions, we performed stimulated emission depletion (STED) microscopy to observe high-resolution sub-compartmental spatial distribution. In *Fbxl21* KO cells, we found a pronounced accumulation of cytoplasmic TDP-43 puncta that co-localized with G3BP1-positive stress granules, whereas TDP-43 remained primarily nuclear in control cells under basal conditions (Fig. 6C). Upon heat shock, the levels of cytoplasmic TDP-43 puncta and G3BP1-positive granules were further elevated in *Fbxl21* KO compared to control cells, with substantial co-localization observed (Fig. 6C). Together, the FBXL21-DNAJB6 axis regulates TDP-43 inclusions and stress granule formation, playing an important role in maintaining proteostasis under cellular stress.

### FBXL21 regulates exercise timing-dependent stress response in skeletal muscle

Timed exercise is an emerging paradigm of circadian skeletal muscle function with a significant impact on metabolism, gene expression, and stress response pathways^58, 59^. To investigate the *in vivo* role of FBXL21-DNAJB6 in modulating stress granule formation in skeletal muscle, we subjected mice to acute treadmill exercise at ZT6 (daytime, the naturally inactive phase for mice) or ZT18 (nighttime) (Fig. 7A). Following exercise, both WT and *Psttm* mice exhibited a significant increase in rectal temperature, confirming effective exercise (Fig. EV5A). DNAJB6 and FBXL21 displayed a clear diurnal pattern in WT mice, with higher expression at ZT18 than ZT6 (Fig. EV5B). DNAJB6 was significantly enriched by exercise regardless of timing, while FBXL21 levels were also elevated with exercise but retained their diurnal pattern (Fig. EV5B). Similarly, *Psttm* mice showed a strong increase in DNAJB6 expression with exercise, and their DNAJB6 levels were consistently higher than those in WT mice (Fig. EV5B). Because exercise imposes transient proteotoxic stress on skeletal muscle, we investigated the impact of timed exercise on the expression of the stress granule markers G3BP1 and FUS. Interestingly, both proteins displayed a diurnal pattern in WT and *Psttm* mice, with greater accumulation observed at ZT6 compared to ZT18 (Fig. 7B,C). Overall, G3BP1 and FUS levels were highly elevated in *Psttm* mice compared to control WT mice at both times (Fig. 7B,C). Acute exercise at both daytime and nighttime further increased G3BP1 and FUS levels in the muscles of *Psttm* mice relative to WT controls (Fig. 7B,C). To determine whether FBXL21 restoration could rescue these protein levels, we delivered muscle-specific AAV-CK6-Fbxl21-HA to WT and *Psttm* mice (FBXL21 expression was confirmed in Fig. EV5C). Strikingly, the daytime exercise–induced increases in G3BP1 and FUS levels were normalized by muscle-specific AAV-CK6-Fbxl21-HA expression in the gastrocnemius in both WT and *Psttm* mice (Fig. 7B,C). These results suggest that the FBXL21-DNAJB6 axis in *Psttm* mice is impaired in client protein degradation, aggravating stress granule accumulation.

**Fig 7.**
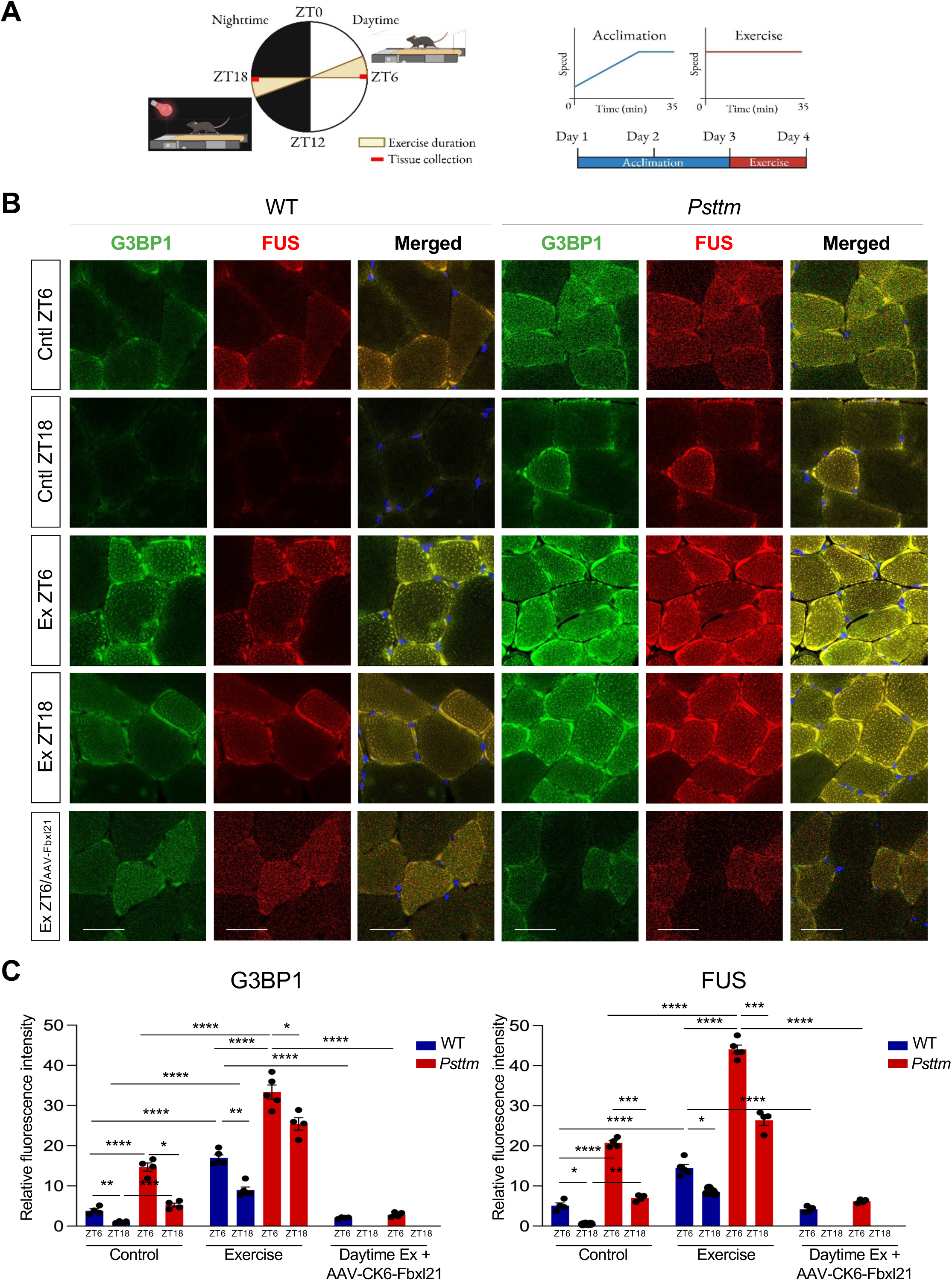
FBXL21 regulates stress response in a timed exercise paradigm. (**A**) Experimental design. After three days of treadmill acclimation at ZT6 and ZT18, WT and *Psttm* mice were subjected on the fourth day to a constant speed of 15 m/min for 35 minutes or until exhaustion. This exercise was performed either during the daytime (ZT6) or nighttime (ZT18). Created in BioRender (https://BioRender.com). (**B**) Immunofluorescent staining of G3BP1 and FUS in gastrocnemius cryosections from 1) WT and *Psttm* mice at ZT6 and ZT18 (control conditions); 2) WT and *Psttm* mice subjected to daytime (ZT6) or nighttime (ZT18) exercise; 3) Skeletal muscle-specific expression of FBXL21 by AAV-CK6-Fbxl21 injection to WT and *Psttm* mice gastrocnemius, and 4 weeks later, mice were subjected to daytime exercise. Imaging was performed using confocal microscopy. Scale bar: 25 μm. (**C**) Quantification of relative fluorescence intensity of G3BP1 (left) and FUS (right). Data are presented as mean ± SEM (n = 4–5 mice/group/time point). No mark indicates not statistically significant. For G3BP1, \**P* = 0.0263 (control *Psttm* ZT6 vs. control *Psttm* ZT18), \**P* = 0.0263 (exercise *Psttm* ZT6 vs. exercise *Psttm* ZT18 \*\**P* = 0.0051 (control WT ZT6 vs. control WT ZT18), \*\*\**P* = 0.0001 (control WT ZT18 vs. control *Psttm* ZT18), and \*\*\*\**P* < 0.0001; Two-way ANOVA indicates statistical differences between groups. For FUS, \**P* = 0.0448 (control WT ZT6 vs. control WT ZT18), \**P* = 0.0448 (exercise WT ZT6 vs. exercise WT ZT18), \*\**P* = 0.0014 (control WT ZT18 vs. control *Psttm* ZT18), \*\*\**P* = 0.0005 (control *Psttm* ZT6 vs. control *Psttm* ZT18), \*\*\**P* = 0.0005 (exercise *Psttm* ZT6 vs. exercise *Psttm* ZT18), and \*\*\*\**P* < 0.0001; Two-way ANOVA indicates statistical differences between groups. N numbers indicate biological replicates for each sample group, and representative data from 3 independent experiments are shown.

TDP-43 and phosphorylated TDP-43 (pTDP-43), the latter known to accumulate in subsarcolemmal protein inclusions in myopathic muscle^60^, were examined by confocal microscopy using gastrocnemius. We found diurnal rhythms of TDP-43 levels in both control and exercised WT mice, with elevated expression in the exercised group (Fig. 8A,B). Interestingly, daytime exercise induced more TDP-43 accumulation than nighttime exercise in WT mice (Fig. 8A,B), possibly due to the lower expression FBXL21-DNAJB6 at ZT6 (Fig. EV5B). In *Psttm* mice without exercise, the diurnal rhythm of TDP-43 was disrupted with elevated levels at both time points (Fig. 8A,B). Importantly, we observed a strong increase in pTDP-43, the pathological marker of inclusions^60^, in exercised *Psttm* mice relative to WT. pTDP-43 accumulation was more pronounced after daytime exercise compared to nighttime exercise in *Psttm* mice, while WT mice showed only mild increase of pTDP-43 in response to daytime exercise (Fig. 8A,B). These results suggest that loss of FBXL21 strongly diminishes DNAJB6 chaperone function under physiological stress conditions. Finally, to determine whether restoring *Fbxl21* expression from *Psttm* mice could mitigate TDP-43 accumulation, we introduced skeletal muscle-specific expression of FBXL21 by AAV-CK6-Fbxl21 delivery to both WT and *Psttm* mice. Muscle-specific expression of *Fbxl21* significantly reduced both TDP-43 and pTDP-43 levels in *Psttm* mice (Fig. 8A,B).

**Fig 8.**
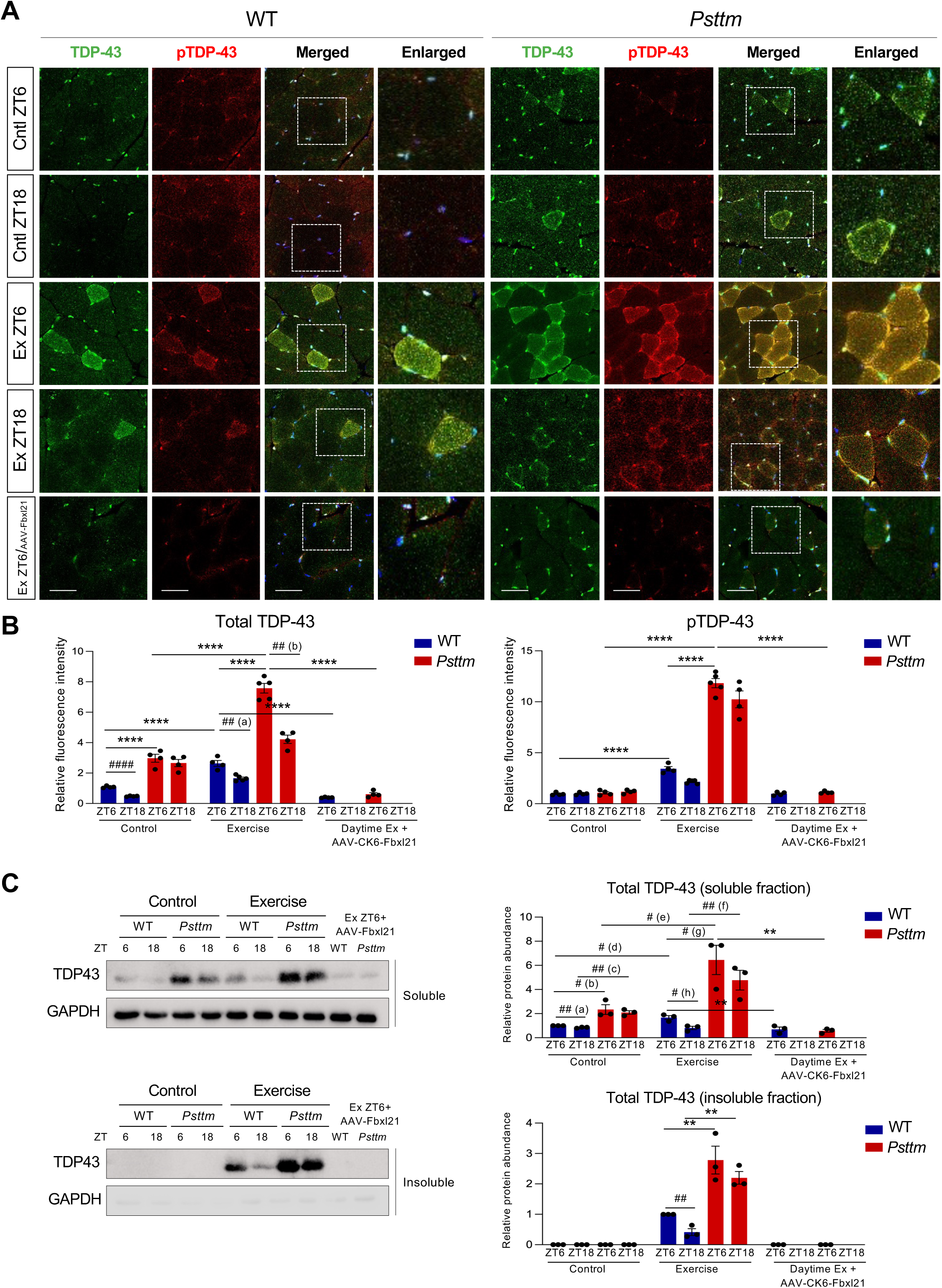
The *Fbxl21* hypomorph *Psttm* mice exhibit aberrant accumulation of TDP-43 and phosphorylated TDP-43 under both normal and timed exercise conditions. (**A**) Immunofluorescence staining of TDP-43 and phosphorylated TDP-43 (pTDP-43) in gastrocnemius cryosections from 1) WT and *Psttm* mice at ZT6 and ZT18 (no exercise, control); 2) WT and *Psttm* mice subjected to daytime or nighttime exercise; 3) WT and *Psttm* mice injected with AAV-CK6-Fbxl21 into the gastrocnemius and subjected to daytime exercise. Imaging was performed using confocal microscopy. Scale bar: 50 μm. (**B**) Quantification of relative fluorescence intensity of total TDP-43 (left) and pTDP-43 (right). Data are presented as mean ± SEM (n = 4–5 mice/group/time point). No mark indicates not statistically significant. \*\*\*\**P* < 0.0001; Two-way ANOVA indicates statistical differences between groups. ^##(a)^*P* = 0.0014, ^##(b)^*P* = 0.0001, and ^####^*P* < 0.0001; Unpaired Student’s *t*-test indicates significant differences between specific comparisons. (**C**) TDP-43 protein levels in the soluble and insoluble fractions from WT and *Psttm* mice subjected to acute exercise. Data are presented as mean ± SEM (n = 3 mice/group/time point). For total TDP-43 (soluble fraction), \*\**P* = 0.0066 (Daytime Ex WT vs. Daytime Ex WT+ AAV-CK6-Fbxl21) and \*\**P* = 0.0066 (Daytime Ex *Psttm* vs. Daytime Ex *Psttm* + AAV-CK6-Fbxl21); Two-way ANOVA indicates statistical differences between groups. ^##(a)^*P* =0.0063, ^#(b)^*P* =0.0289, ^##(c)^*P* =0.0012, ^#(d)^*P* =0.0141, ^#(e)^*P* =0.0324, ^##(f)^*P* =0.0087, ^#(g)^*P* =0.0175, and ^#(h)^*P* =0.0157; Student’s *t*-test indicates significant differences between specific comparisons. For total TDP-43 (insoluble fraction), \*\**P* = 0.0051 (exercise WT ZT6 vs. exercise *Psttm* ZT6), \*\**P* = 0.0052 (exercise WT ZT18 vs. exercise *Psttm* ZT18); One-way ANOVA indicates statistical differences between groups. ^##^*P* = 0.0067; Unpaired Student’s *t*-test indicates significant differences between specific comparisons. N numbers indicate biological replicates for each sample group and representative data from 3 independent experiments are shown.

Finally, we performed detergent-based soluble and insoluble fractionation of gastrocnemius muscle lysates from WT and *Psttm* mice at ZT6 and ZT18, followed by immunoblotting for TDP-43. Consistent with the immunofluorescence staining results, soluble TDP-43 exhibited statistically significant diurnal rhythms without exercise only in WT mice, while *Psttm* mice showed significantly elevated levels at both time points. Daytime exercise increased TDP-43 levels in both WT and *Psttm* mice, again with higher levels in the latter. In the insoluble fractions, TDP-43 was detected only after acute exercise, with increased amounts in *Psttm* mice compared with WT mice (Fig. 8C). Interestingly, Muscle-specific expression of *Fbxl21* significantly reduced the accumulation of soluble and insoluble TDP-43 (Fig. 8C). These results suggest that FBXL21 may regulate stress-induced accumulation of insoluble TDP-43 in skeletal muscle.

## Discussion

Circadian regulation of proteostasis, including protein folding, modification, and degradation, is critical for physiological fitness. The circadian E3 ligase FBXL21 governs target protein oscillation by ubiquitination-mediated protein degradation^18–21^. Our current study reveals a novel role of FBXL21 as a chaperone-linked E3 ubiquitin ligase in the diurnal regulation of skeletal muscle proteostasis. We demonstrate that FBXL21 regulates the degradation and diurnal rhythms of DNAJB6, a HSP40 family co-chaperone, as well as its client proteins such as Desmin, via ubiquitination-mediated degradation. In contrast to WT, disease-causing DNAJB6 mutations confer protection from FBXL21-mediated degradation. Whereas HSP70 also interacts with FBXL21 as a molecular chaperone to facilitate FBXL21 folding and E3 ligase activity, it is not itself a target of FBXL21-mediated degradation. Furthermore, our data suggest that FBXL21 functions independently from the HSP70-linked E3 ligase CHIP in DNAJB6 client protein degradation, underscoring the selectivity of FBXL21 toward DNAJB6. Analogous to CHIP which targets dysfunctional HSP70 for ubiquitination and degradation to maintain proteostasis^43^, FBXL21 regulates DNAJB6 turnover, mitigating the accumulation of damaged or misfolded proteins. On the other hand, Desmin-positive inclusion bodies and TDP-43 inclusions were observed in the *Fbxl21* hypomorph *Psttm* mice, indicative of proteotoxic muscle stress similar to those observed with DNAJB6 G/F domain mutants^54, 57^. Using timed exercise as a physiological stressor, we further found that FBXL21 contributes to the diurnal regulation of stress responses in skeletal muscle. In *Fbxl21* hypomorphic *Psttm* mice, levels of proteostatic markers such as G3BP1, FUS, and TDP-43 remained higher at ZT6 than at ZT18, yet elevated at both time points when compared to WT mice, suggesting an altered diurnal rhythm rather than complete loss of rhythmicity. These abnormalities were mitigated by the restoration of FBXL21 expression. Together, our results position FBXL21 as a key node linking molecular chaperones, the ubiquitin-proteasome system and the circadian machinery to maintain muscle proteostasis.

In protein quality control, misfolded proteins are targeted for proteasomal degradation via ubiquitination, mediated by specific chaperone–E3 ligase mechanisms. A well-characterized example in mammals is the HSP70-CHIP pathway^28, 43, 61^. Whereas HSP70-CHIP is required for disposal of terminally misfolded proteins, recent findings in yeast suggest that distinct pathways may be involved for processing misfolded proteins. In yeast, the Sis1 (Hsp40)-Doa10 (E3 ligase) complex is required for ubiquitination of its misfolded client protein Vma12-DegAB^62, 63^. Likewise, we show that DNAJB6, in addition to its established role of transferring structurally abnormal client proteins to HSP70-CHIP, plays a critical role in misfold protein degradation by recruiting FBXL21, highlighting a regulatory function in proteostasis by selectively guiding aberrant proteins toward degradation^62^.

Humans have more than 600 E3 ubiquitin ligases, each with specificity for a distinct subset of target proteins^64^. In the skeletal muscle, FBXL21^20, 21^, Atrogin-1, and MuRF1 are known to show circadian transcript expression and mediate circadian time-dependent protein degradation. Overall, 48 out of 941 rhythmically expressed genes (5%) in the gastrocnemius muscle encode circadian E3 ubiquitin ligases^8^, suggesting an important role of circadian E3 ligases in skeletal muscle proteostasis. E3 ligases can directly target circadian substrates. For example, Atrogin-1 (FBXO32) and MuRF1 degrade transcription factors or structural proteins in a time-dependent manner, aligning muscle protein turnover with activity and rest cycles^65^. E3 ligases can also function by mediating the degradation of chaperone-bound client proteins. As mentioned above, CHIP (STUB1) associates with HSP70/HSP90 to eliminate misfolded or excess proteins. This chaperone-client degradation pathway may be more pervasive than originally thought in terms of target proteins^33, 56, 66^. Notably, FBXL21 appears to function in both modes. While our previous work revealed that FBXL21 controls the stability and circadian rhythms of the Z-line proteins TCAP and MYOZ1^20, 21^, the current study demonstrates its activity in regulating the degradation of chaperone client proteins. This dual functionality encapsulates the pivotal roles of FBXL21: it not only regulates core clock components (CRY1, CRY2)^18, 19^ and other specific substrate targets ^20, 21^, but also controls broader proteostasis networks through chaperone-mediated pathways.

Emerging evidence highlights a dynamic and reciprocal interplay between circadian clock components and molecular chaperones such as HSP70 and HSP90. Core clock proteins such as PER2 and BMAL1 are directly regulated by chaperones, which impacts their stability, localization, and function. Specifically, PER2 interacts with both HSP70 and HSP90 to promote cuproptosis^22^ and antitumor immune responses^23^, respectively. HSP90 also stabilizes BMAL1, and its inhibition leads to reduced circadian gene expression in cultured cells^24^. Furthermore, PER2 regulates chaperone-mediated autophagy (CMA) which reciprocally influences circadian protein turnover, suggesting a bidirectional feedback loop^67^. In plants, the circadian F-box protein ZEITLUPE (ZTL) is a client of HSP90^68^, with the co-chaperone GIGANTEA facilitating its maturation^69^; ZTL in turn contributes to thermo-responsive proteostasis^70^. Analogous to the ZTL–HSP90 interaction, the FBXL21–HSP70 axis functions via an evolutionarily conserved mechanism where a circadian F-box protein requires the chaperone–co-chaperone system for stability and concomitantly cooperates to regulate client protein degradation.

DNAJB6, a member of the HSP40 family of co-chaperones, partners with HSP70 to suppress aggregation of misfolded or aggregation-prone proteins. Dominant mutations in DNAJB6 – particularly within its glycine/phenylalanine (G/F)-rich domain – cause limb-girdle muscular dystrophy type 1D (LGMD1D), a progressive skeletal muscle disease marked by myofibrillar disruption, rimmed vacuoles, and protein inclusions^36, 39, 54, 56, 71^. In skeletal muscle, amyloidogenesis, the misfolding and aggregation of proteins into insoluble fibrils, contributes to degenerative myopathies when protein quality control fails; DNAJB6 chaperone function is a key suppressor of this process ^35, 51, 55, 60, 71^. Mutations in DNAJB6 compromise this protective function, leading to accumulation of toxic protein inclusions and subsequently muscle fiber degeneration^33, 34, 36, 56^. For example, defective DNAJB6 chaperone activity has been shown to cause aggregation of RNA-binding proteins and Z-disk pathology^54, 56, 71^. Previous studies showed that the *Fbxl21* hypomorphic *Psttm* mice exhibit impaired degradation of sarcomeric substrates and reduced muscle performance, including decreased running distance at nighttime and grip strength^20^. Of note, accumulating evidence indicates that dysregulated proteostasis and subsequent accumulation of aggregation-prone proteins, such as TDP-43 and Desmin, can adversely affect muscle structure and performance^50, 51, 55, 60, 72–76^. In line with these findings, we show that disruption of the FBXL21–DNAJB6 axis in *Psttm* mice perturbs diurnal proteostasis and aggravates accumulation of aggregation-prone proteins, which may contribute to skeletal muscle dysfunction^20^. Finally, our current work shows that the circadian FBXL21-DNAJB6 axis targets misfolded chaperone clients for proteasomal degradation. When DNAJB6 binds to client proteins, FBXL21 shifts its activity from ubiquitinating DNAJB6 itself to DNAJB6-bound client proteins, suggesting coordinated degradation to resolve misfolded or aggregation-prone substrates. Failure of this degradation axis due to a loss of FBXL21 exacerbates toxic protein accumulation and impairs stress response to timed exercise. Together, these observations highlight FBXL21–DNAJB6 as a new chaperone-associated protein ubiquitination machinery to maintain skeletal muscle proteostasis.

In conclusion, our findings elucidate a novel regulatory mechanism in which FBXL21 cooperates with DNAJB6 to facilitate client protein degradation. Loss of FBXL21 disrupts this axis, leading to pathological protein accumulation and aggregation. The FBXL21–DNAJB6 axis may represent a critical quality control pathway for muscle homeostasis and a potential target for therapeutic intervention against protein aggregation disorders.

## Materials & Methods

### Animal Studies and Ethics Statement

C57BL/6J mice (Stock #000664; Jackson Laboratory, Bar Harbor, ME, USA) were purchased, and *Psttm* and wild-type (WT) littermates were bred in-house. Unless otherwise specified, all animals were maintained under a 12 h light/12 h dark cycle (LD 12:12). Male mice were used for this study. All animal husbandry and experiments, including considerations of sample size, randomization and blinding, were approved by the Center for Laboratory Animal Medicine and Care (CLAMC) at The University of Texas Health Science Center at Houston (UTHealth Houston), under the protocol #AWC-23-0038.

### Yeast 2-hybrid (Y2H) screen

A yeast two-hybrid (Y2H) screen was performed using FBXL21 as bait against a human skeletal muscle cDNA library in the AH109 yeast strain, as previously described ^20^. The screen identified over 30 interacting proteins. Among the top candidates, DNAJB6 was selected for further validation.

### Mass spectrometry (MS) analysis

To detect the interactome of FBXL21, we performed immunoaffinity purification followed by nanoLC-MS/MS analysis, based on a previously reported method with minor modifications. Briefly, the affinity-purified proteins, along with their interacting partners, were digested directly on the beads overnight using Trypsin/Lys-C (Thermo Scientific™, A40007). The resulting peptides were then purified and desalted using in-house–prepared STAGE tips containing 2 mg of C18 material (3 µm, Dr. Maisch GmbH, Germany), following established procedures^77^. After desalting, peptides were dried under vacuum and resuspended for LC–MS analysis on an nLC 1000 system coupled to an Orbitrap Fusion mass spectrometer (Thermo Scientific) equipped with an electrospray ionization source. Mass spectrometry data were acquired using a combined method of data-dependent acquisition (DDA) for unbiased peptide profiling. Data analysis was conducted using Proteome Discoverer 2.1 (Thermo Fisher) with the Mascot algorithm (Mascot 2.4, Matrix Science).

### Cell Culture, Transfection, and Differentiation

HEK293T (ATCC CRL-3216) and C2C12 (ATCC CRL-1772) cells were cultured in DMEM supplemented with 10% fetal bovine serum (FBS; GenDEPOT, Houston, TX, USA). For immunoprecipitations, 2 × 10⁶ cells were seeded in 60-mm dishes 24 h prior to transfection, and plasmids were introduced using iMFectin (GenDEPOT) according to the manufacturer’s protocol. C2C12 cells were cultured in DMEM containing 10% FBS and penicillin/streptomycin until reaching 80–90% confluence. Differentiation was induced by switching to DMEM supplemented with 2% horse serum and penicillin/streptomycin.

### siRNA Transfection

HEK293T cells were seeded at a density of 2 × 10⁵ cells per well in 6-well plates and allowed to adhere overnight. For gene knockdown experiments, cells were transfected with 50 nM of validated MISSION siRNAs targeting HSP70 (HSPA1A; SASI_HS01_00051559, Sigma-Aldrich, St. Louis, MO, USA), CHIP (Stub1; SASI_HS01_00183572, Sigma-Aldrich, St. Louis, MO, USA), DNAJB6 (DNAJB6; SASI_HS01_00212271) or a scrambled control siRNA (Dharmacon, Lafayette, CO, USA; or Ambion, Austin, TX, USA) using Lipofectamine RNAiMAX (Thermo Fisher Scientific, Waltham, MA, USA) according to the manufacturer’s instructions. In brief, siRNA and RNAiMAX were diluted in Opti-MEM (Thermo Fisher Scientific), mixed, and incubated for 15 minutes at room temperature to allow complex assembly. The transfection complexes were then added dropwise to cells maintained in antibiotic-free DMEM supplemented with 10% FBS. Knockdown efficiency was confirmed by immunoblotting.

### RNA extraction and real-time qPCR

Total RNA was extracted from frozen gastrocnemius tissues using PureXtract RNAsol RNA isolation solution (GenDEPOT) as previously described^21^. cDNA was synthesized from 2 µg of total RNA, and PCR amplification was performed using amfiSure qGreen Q-PCR Master Mix (GenDEPOT). Relative mRNA abundance was obtained by a comparative Ct method (ΔΔCt method)^78^. GAPDH was used as a loading control. The primer sequences used were as follows: *Dnajb6*-F: GAAGTTGAAGAAGATGGGCAGT, *Dnajb6*-R: CTCTCCGCTGGCATTCTTCT, *Gapdh*-F: CAAGGTCATCCATGACAACTTTG, and *Gapdh*-R: GGCCATCCACAGTCTTCTGG.

### Detergent-soluble and insoluble fractionation

Gastrocnemius tissues from WT and *Psttm* mice were homogenized in 0.5% Triton X-100 lysis buffer supplemented with protease and phosphatase inhibitors. The homogenates were centrifuged at 13,000 rpm for 15 min at 4 °C to pellet detergent-insoluble fractions. The supernatant is collected as the soluble fraction. For the insoluble fraction, the remaining pellets were resuspended in RIPA buffer containing 6M urea and 2% SDS, sonicated for 15 seconds, and centrifuged at 14,000 rpm for 10 minutes at 4 °C. The resulting supernatant was collected as the insoluble fraction. GAPDH was used as a positive control of the soluble fraction. Total lysates (Figs. 1D, EV3C) were prepared using RIPA buffer. For fractionation experiments (Fig. EV1E / RL Fig. S1H, Fig. EV3D/RL-Fig8), gastrocnemius muscles were extracted with Triton X-100 to obtain the soluble fraction, followed by urea/SDS buffer to extract the detergent-insoluble fraction.

### Immunoblotting, Immunoprecipitation, and immunofluorescence staining

Immunoblotting, immunoprecipitation, and immunofluorescence staining were performed as previously described ^18, 20, 21^. For protein degradation assays, 2 × 10⁵ HEK293T cells were seeded in 12-well plates and transfected with expression constructs encoding pCMV10-3XFlag-Dnajb6b, pCMV10-3XFlag-Desmin, V5-HSP70, and pCMV10-3XFlag-Fbxl21 constructs. After 36 hours, cells were treated with cycloheximide (CHX, 100 μg/mL), harvested at the indicated time points, and protein half-lives were determined using nonlinear one-phase decay analysis (GraphPad Prism). For immunoblotting, detection of ectopically expressed proteins was performed using anti-Flag-HRP (Sigma-Aldrich), anti-HA (Roche), and anti-V5 (Thermo Fisher Scientific) antibodies. Endogenous proteins in skeletal muscle and C2C12 cells were detected using respective antibodies, including DNAJB6 (ProteinTech), Desmin (Abcam), HSP70 (ProteinTech), and GAPDH (Abclonal Biotechnology). For polyQ23htt-EGFP and polyQ74htt-EGFP staining, control and *Fbxl21* KO C2C12 cells were transfected with polyQ23htt-EGFP (Addgene #40261) and polyQ74htt-EGFP (Addgene #40262) using Lipofectamine 3000 (Thermo Fisher Scientific). Twenty-four hours post-transfection, cells were subjected to one of the following conditions: no heat shock (control), heat shock (HS) at 42 °C, or HS followed by a 2-hour recovery. The cells were then fixed with 4% paraformaldehyde and imaged for GFP signal using confocal microscopy (Leica Microsystems, Mannheim, Germany). Aggregated polyQ74htt-EGFP was quantified by measuring relative aggregate area (%) according to the published method^79–81^. Specifically, the total aggregate area (%) per cell was measured and normalized to the corresponding total cell area, then multiplied by 100 to obtain the percentage of aggregate burden. For *in vitro* quantification, we counted more than 100 cells per group with uniform thresholding criteria applied across all groups. For immunofluorescence staining of endogenous proteins, antibodies against DNAJB6 (ProteinTech), Desmin (ABclonal Biotechnology, China; and Abcam, Cambridge, MA, USA), TDP-43 (ProteinTech), and phospho-TDP-43 (ProteinTech), G3BP1 (Invitrogen), and FUS (Invitrogen) were used. Anti-Fbxl21 antibodies were custom-generated in rabbits (Cocalico Biologicals, Reamstown, PA, USA) and affinity-purified against the immunizing peptide secondary antibodies for confocal imaging were goat anti-mouse Alexa Fluor 488, goat anti-mouse Alexa Fluor 568, goat anti-rabbit Alexa Fluor 488, and goat anti-rabbit Alexa fluor 568 (all from Invitrogen). For STED imaging, secondary antibodies were goat anti-mouse STAR ORANGE and goat anti-rabbit STAR RED (Abberior). Images were acquired using a STELLARIS confocal microscope (Leica Microsystems) or STED microscope (Abberior Instruments, Göttingen, Germany). Ubiquitination assays were conducted as previously described^18, 20, 21^).

### *In vivo* muscle immunofluorescence staining and quantification

Skeletal muscle samples were embedded in O.C.T. compound (Tissue-Tek), snap-frozen in liquid nitrogen-cooled isopentane, stored at −80 °C, and sectioned at 20 μm thickness using a cryostat (Leica Biosystems, Germany). The sections were pre-treated with heat-mediated antigen retrieval using 10 mM citrate buffer (pH 6.0). After blocking with 1% BSA, 0.2% Triton X-100, and 5% goat serum in PBS, sections were incubated overnight at 4°C with primary antibodies of DNAJB6 (1:500), TDP-43 (1:1000), and pTDP-43 (1:500) (all from Proteintech); G3BP1 (1:400) and FUS (1:500) (Invitrogen). Sections were then incubated with secondary antibodies (1:1000) for 1 hour at room temperature. Images were obtained using a STELLARIS confocal microscope (Leica Microsystems) with identical acquisition settings (laser power, gain, and exposure) across all samples. DNAJB6 and Desmin inclusions were defined as clear puncta or inclusion-like cytoplasmic structures that are distinguishable from the regular diffuse staining pattern as previously reported^33, 82–84^. For quantification of DNAJB6 and desmin inclusions, comparable random regions of the gastrocnemius (three to five random fields per section) were selected across all samples, and the number of inclusions was quantified per approximately 500 fibers per sample, and % of fibers containing inclusions was calculated. For measurement of relative fluorescence intensity, comparable random regions of the gastrocnemius (three to five random fields) were selected and imaged using a STELLARIS confocal microscope (Leica Microsystems) with identical acquisition settings (including laser power, gain, and exposure) across all samples. Relative fluorescence intensity was measured as previously described^85–87^. All images were converted into 8-bit grayscale. The “threshold” function was used to perform binary segmentation. The mean fluorescence intensity was quantified within defined regions of interest (ROIs) using uniform acquisition settings and consistent thresholding parameters applied across all groups. Signal intensity was normalized to background to ensure comparability. A total of more than 1,500 muscle fibers per mouse were evaluated for all figures.

### Vector design and muscle-specific delivery of AAV-Fbxl21

Mouse Fbxl21 cDNA was subcloned into pAAV-CK6 with a C-terminus HA tag ^88^. The AAV9-CK6-Fbxl21-HA virus (serotype 9) was produced. Eight-week-old WT and *Psttm* mice were injected with 1 × 10^12^ genome copies (GC) of AAV into the gastrocnemius muscle. Four weeks post-injection, the mice were subjected to the exercise paradigm.

### Acute exercise paradigm

The treadmill exercise test was performed as previously described ^89^ with minor modifications. WT and *Psttm* mice at 5 months old were acclimated to the Exer-3R treadmill (Columbus Equipment) by running at gradually increasing speeds up to 15 m/ min for 35 minutes per day over three consecutive days. On the fourth day, the mice were subjected to a constant speed of 15 m/min for 35 minutes or until exhaustion. Immediately following the exercise, the mice were euthanized, and calf muscle tissue was collected. The assay was performed at two circadian time points, ZT6 and ZT18, using different cohorts of mice for each time point.

### Quantification and Statistical Analysis

Data are presented as mean ± SEM. All the n numbers indicate biological replicates for each sample group. For *in vivo* studies, n corresponds to mouse number. In cell studies, n numbers refer to independent cultures for each experiment, and each experiment has been repeated to ensure reproducibility. Specifically, in each independent experiment, three or more biological replicates were included, corresponding to separate wells undergoing parallel transfection and treatment, sample preparation, and western blot analysis. The experiment was repeated for a total of three times or more. Similar results were obtained from the independent experiments, and representative data are shown in the figures. Statistical analyses were conducted using GraphPad Prism (GraphPad Software, Inc.). Comparisons were made using Student’s *t*-test or one-way/two-way ANOVA, followed by Tukey’s post hoc test. Statistical significance was defined as *P* < 0.05.

## Acknowledgements

We thank Kaori Ono for technical support. We thank the Center for Advanced Microscopy of the UTHealth McGovern Medical School for technical assistance. This work is supported by NIH/NIGMS (R35GM145232) and The Welch Foundation (AU-2127-20220331) to S.-H.Y, NIH/NIA (R01AG089967, R01AG065984) to Z.C., (R01AR079220) to K.A.E., and (R01AI158694) to K.E-M (R01A1158694).

## Conflict of Interest

The authors declare no conflict of interest.

## Figure legends for EV

**Figure EV1. Identification of DNAJB6 as a novel target of FBXL21.**

(A) Yeast two-hybrid (Y2H) assay demonstrating the interaction between FBXL21 and DNAJB6. As a positive control, yeast cells were co-transformed with pGBKT7-p53 (bait) and pGADT7-SV40 (prey), expressing the Gal4 DNA-binding domain (BD) fused to murine p53 and the Gal4 activation domain (AD) fused to the SV40 large T antigen, respectively. (**B–C**) Coimmunoprecipitation assays showing the interaction between FBXL21 and human DNAJB6a or DNAJB6b. (**D**) mRNA expression of *Dnajb6* in WT and *Psttm* mice. Data are presented as mean ± SEM (n = 5/group/time point). No mark indicates not statistically significant. (**E**) Immunoblot of DNAJB6 in soluble and insoluble fractions from WT and *Psttm* mice. GAPDH was used as a positive control for the soluble fraction. Data are presented as mean ± SEM (n = 3 mice/group/time point). \*\*\**P* = 0.0002 and \*\*\*\**P* < 0.0001; Two-way ANOVA with Tukey’s multiple comparisons test, comparing between WT and *Psttm* mice or between time points. ^###^*P* = 0.0002 and ^####^*P* < 0.0001; Unpaired Student’s t-test indicates significant differences between specific comparisons. (**F**) Immunoblot analysis of FBXL21 expression in Cntl and *Fbxl21* KO C2C12 cells. Data are presented as mean ± SEM (n = 3). \**P* = 0.0363, Student’s t-test, indicating a significant difference between groups. (**G**) Immunofluorescence staining of endogenous FBXL21 in Cntl and *Fbxl21* KO C2C12 cells by confocal microscopy. Scale bar: 20 μm. (**H**) Negative control image (secondary antibody only) for Fig. 1F. N numbers indicate biological replicates for each sample group, and representative data from three independent experiments are shown.

**Figure EV2. FBXL21 regulates degradation and ubiquitination of human DNAJB6 isoforms.**

(**A–B**) Cycloheximide (CHX) chase assays were performed to assess the protein stability of human DNAJB6a (A) and DNAJB6b (**B**) in 293T cells. Protein half-lives were quantified. Half-lives: hDnajb6a, 7.05 h; hDnajb6a/Fbxl21, 2.82 h; hDnajb6b, >24 h; hDnajb6b/Fbxl21. Data are presented as mean ± SEM (n = 3). (**C–D**) FBXL21 promotes the ubiquitination of human DNAJB6 isoforms. Flag-tagged DNAJB6a (**C**) or DNAJB6b (**D**) was co-transfected with untagged FBXL21 and HA-tagged ubiquitin (HA-Ub), followed by ubiquitination assays. (**E**) ΔFBXL21 abolished its ability to ubiquitinate DNAJB6b in 293T cells. Flag-tagged mDNAJB6b was co-transfected with either wild-type FBXL21 or ΔFBXL21 with HA-tagged ubiquitin (HA-Ub), followed by ubiquitination assays. N numbers indicate biological replicates for each sample group, and representative data from three independent experiments are shown.

**Figure EV3. FBXL21 regulates desmin proteostasis and polyQ74htt aggregation without altering HSP70 expression.** (**A**) Immunoblot analysis of HSP70 protein levels in wild-type (WT) and *Psttm* mice collected at Zeitgeber time 6 (ZT6; daytime) and ZT18 (nighttime). GAPDH was used as a loading control. Data are presented as mean ± SEM (n = 3/group/time point). No mark indicates not statistically significant. *P* = 0.7027 (WT ZT6 vs. *Psttm* ZT6) and *P* = 0.8803 (WT ZT18 vs. *Psttm* ZT18); Two-way ANOVA with Tukey’s multiple comparisons test. (**B**) Immunoblot analysis of HSP70 protein levels in control and *Fbxl21* KO C2C1 cells. Data are presented as mean ± SEM (n = 3). No mark indicates not statistically significant. *P* = 0.0816; Unpaired Student’s t-test. (**C**) Immunoblot analysis of Desmin protein levels in WT and *Psttm* mice. GAPDH was used as a loading control. Data are presented as mean ± SEM (n = 3–4). \*\**P* = 0.0059 *and* \*\*\**P* = 0.0004; two-way ANOVA indicates statistical differences between genotypes (WT vs. *Psttm*). (**D**) Immunoblot of Desmin in soluble and insoluble fractions from WT and *Psttm* mice. GAPDH was used as a positive control for the soluble fraction. Data are presented as mean ± SEM (n = 3). \*\*\**P* = 0.0001 and \*\*\*\**P* < 0.0001; Two-way ANOVA with Tukey’s multiple comparisons test, comparing between WT and *Psttm* mice or between time points. (**E**) Immunoblot analysis of Desmin protein levels in Cntl and *Fbxl21* KO C2C12 cells. GAPDH was used as a loading control. Data are presented as mean ± SEM (n = 3). **\*\*\*\****P* < 0.0001; Unpaired Student’s t-test. (**F**) Confocal microscopy analysis in Cntl and *Fbxl21* KO C2C12 cells expressing polyQ23htt-EGFP and polyQ74htt-EGFP. PolyQ23htt-EGFP was used as a negative control, and no aggregates were detected in the groups. The relative Q74 aggregate area (%) was quantified using ImageJ. Data are presented as mean ± SEM (n=100 cells/group). **\****P* = 0.0253 (WT HS vs. WT HS recovery), \*\**P* = 0.0023 (WT no HS vs. WT HS), and \*\*\*\**P* < 0.0001; One-way ANOVA showing significant differences between groups. **^####^***P* < 0.0001; Unpaired Student’s *t*-test. N numbers indicate biological replicates for each sample group, and representative data from three independent experiments are shown.

**Figure EV4. DNAJB6 and FBXL21 colocalize in stress granules. Left panel:** Confocal microscopy of HEK293T cells co-expressing either Flag-DNAJB6b and G3BP1-RFP or Flag-FBXL21 and G3BP1-RFP. Scale bar: 20 μm. **Right panels**: Line profiles showing the mean fluorescence intensities analyzed by using Prism. Representative images are shown from three independent experiments are shown. Scale bar: 20 μm.

**Figure EV5. Exercise alters DNAJB6 and FBXL21 expression levels in WT and *Psttm* mice in a time-dependent manner.** (**A**) Rectal temperature measured using a rectal probe (ThermoWorks) at different time points. Data are presented as mean ± SEM (n = 5 mice/group/time point). No mark indicates not statistically significant. \*\**P* = 0.0017 and \*\*\*\**P* < 0.0001, Two-way ANOVA with Tukey’s multiple comparisons test, comparing between groups. (A) Immunofluorescence staining of DNAJB6 and FBXL21 in gastrocnemius muscle cross-sections from wild-type (WT) and *Psttm* mice under control (non-exercise) conditions. Scale bar: 20 μm. Lower left panel: Quantification of DNAJB6 fluorescence intensity in gastrocnemius muscle cross-sections from WT and *Psttm* mice under control and acute exercise conditions. Lower right panel: Quantification of FBXL21 fluorescence intensity in gastrocnemius muscle cross-sections from WT and *Psttm* mice under control and acute exercise conditions. Data are presented as mean ± SEM (n = 3/group/time point). For DNAJB6, \**P* = 0.0483 (control WT ZT18 vs. control *Psttm* ZT18), \**P* = 0.0131 (exercise WT ZT6 vs. exercise *Psttm* ZT6), \*\**P* = 0.0061 (exercise WT ZT18 vs. exercise *Psttm* ZT18), \*\*\**P =* 0.0015 (control WT ZT6 vs. exercise WT ZT6), \*\*\**P* = 0.0001 (exercise WT ZT18 vs. exercise *Psttm* ZT18), and \*\*\*\**P* < 0.0001; Two-way ANOVA with Tukey’s multiple comparisons test. ^##^*P* = 0.0028 and ^###^*P* = 0.0010; Unpaired Student’s *t*-test. For FBXL21, \**P* = 0.0176 and \*\*\*\**P* < 0.0001; Two-way ANOVA with Tukey’s multiple comparisons test. (**C**) Immunoblot assay for HA detection in gastrocnemius tissues of WT and *Psttm* mice subjected to daytime exercise and AAV-CK6-Fbxl21-HA injection.

## Notes

### Competing Interest Statement

The authors have declared no competing interest.

