## Supplemental materials for "FBXL21 regulates diurnal proteostasis and stress response by targeting DNAJB6 and client proteins"

### Fig. EV1

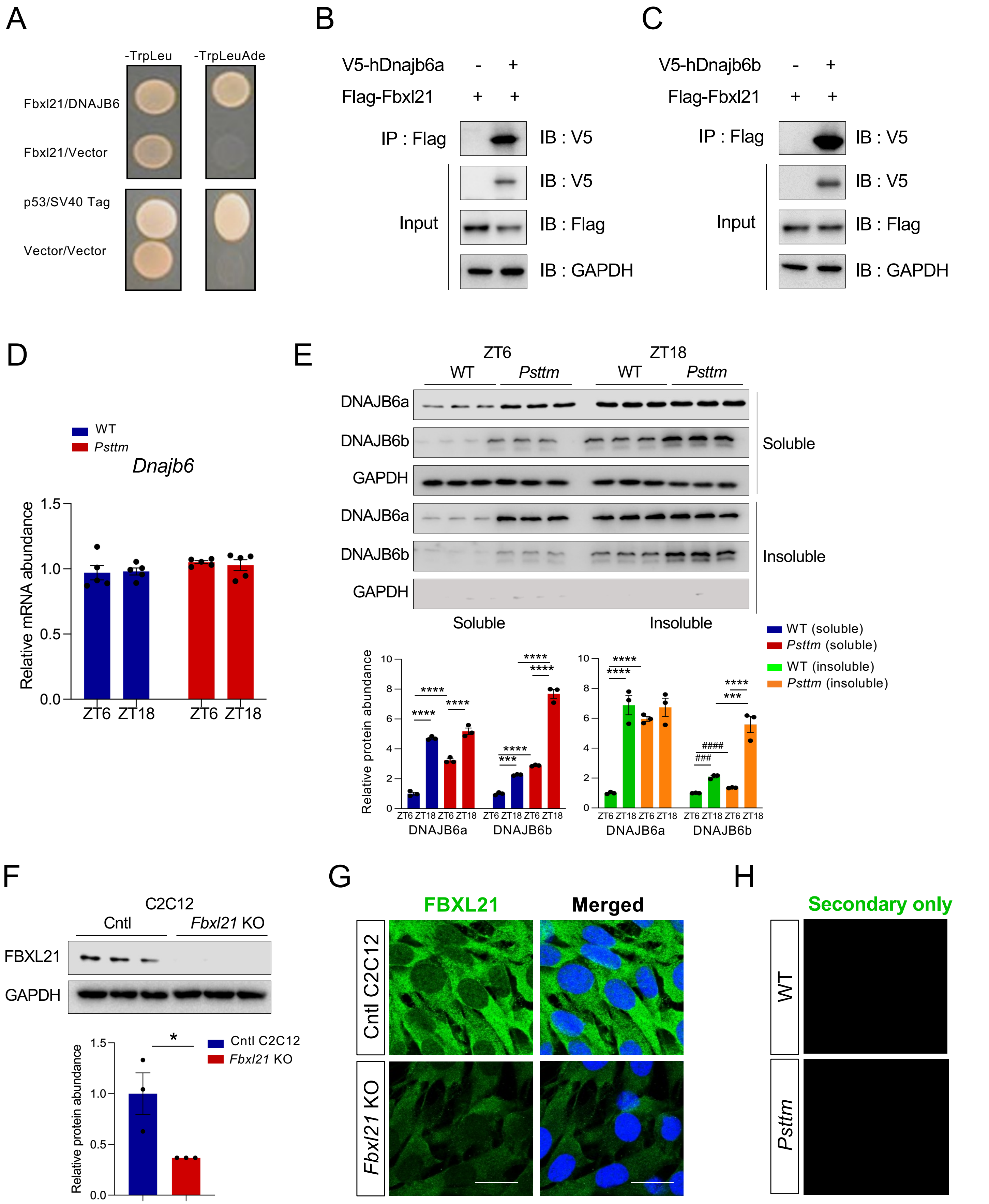
A
-TrpLeu
-TrpLeuAde
Fbxl21/DNAJB6
Fbxl21/Vector
p53/SV40 Tag
Vector/Vector
B
V5-hDnajb6a - +
Flag-Fbxl21 + +
IP : Flag
IB : V5
Input
IB : V5
IB : Flag
IB : GAPDH
C
V5-hDnajb6b - +
Flag-Fbxl21 + +
IP : Flag
IB : V5
Input
IB : V5
IB : Flag
IB : GAPDH
D
WT
Psttm
Dnajb6
Relative mRNA abundance
ZT6 ZT18 ZT6 ZT18
E
ZT6
ZT18
WT
Psttm
DNAJB6a
DNAJB6b
GAPDH
Soluble
Insoluble
WT (soluble)
Psttm (soluble)
WT (insoluble)
Psttm (insoluble)
Relative protein abundance
ZT6 ZT18 ZT6 ZT18
DNAJB6a
DNAJB6b
F
C2C12
Cntl
Fbxl21 KO
FBXL21
GAPDH
Cntl C2C12
Fbxl21 KO
Relative protein abundance
G
FBXL21
Merged
Cntl C2C12
Fbxl21 KO
H
Secondary only
WT
Psttm

**Figure EV1. Identification of DNAJB6 as a novel target of FBXL21.**

(**A**) Yeast two-hybrid (Y2H) assay demonstrating the interaction between FBXL21 and DNAJB6. As a positive control, yeast cells were co-transformed with pGBKT7-p53 (bait) and pGADT7-SV40 (prey), expressing the Gal4 DNA-binding domain (BD) fused to murine p53 and the Gal4 activation domain (AD) fused to the SV40 large T antigen, respectively. (**B–C**) Coimmunoprecipitation assays showing the interaction between FBXL21 and human DNAJB6a or DNAJB6b. (**D**) mRNA expression of *Dnajb6* in WT and *Psttm* mice. Data are presented as mean ± SEM (n = 5/group/time point). No mark indicates not statistically significant. (**E**) Immunoblot of DNAJB6 in soluble and insoluble fractions from WT and *Psttm* mice. GAPDH was used as a positive control for the soluble fraction. Data are presented as mean ± SEM (n = 3 mice/group/time point). ****P* = 0.0002 and *****P* < 0.0001; Two-way ANOVA with Tukey's multiple comparisons test, comparing between WT and *Psttm* mice or between time points. $^{\#\#\#}P$ = 0.0002 and $^{\#\#\#\#}P$ < 0.0001; Unpaired Student's t-test indicates significant differences between specific comparisons. (**F**) Immunoblot analysis of FBXL21 expression in Cntl and *Fbxl21* KO C2C12 cells. Data are presented as mean ± SEM (n = 3). **P* = 0.0363, Student's t-test, indicating a significant difference between groups. (**G**) Immunofluorescence staining of endogenous FBXL21 in Cntl and *Fbxl21* KO C2C12 cells by confocal microscopy. Scale bar: 20 μm. (**H**) Negative control image (secondary antibody only) for Fig. 1F. N numbers indicate biological replicates for each sample group, and representative data from three independent experiments are shown.

### Fig. EV2

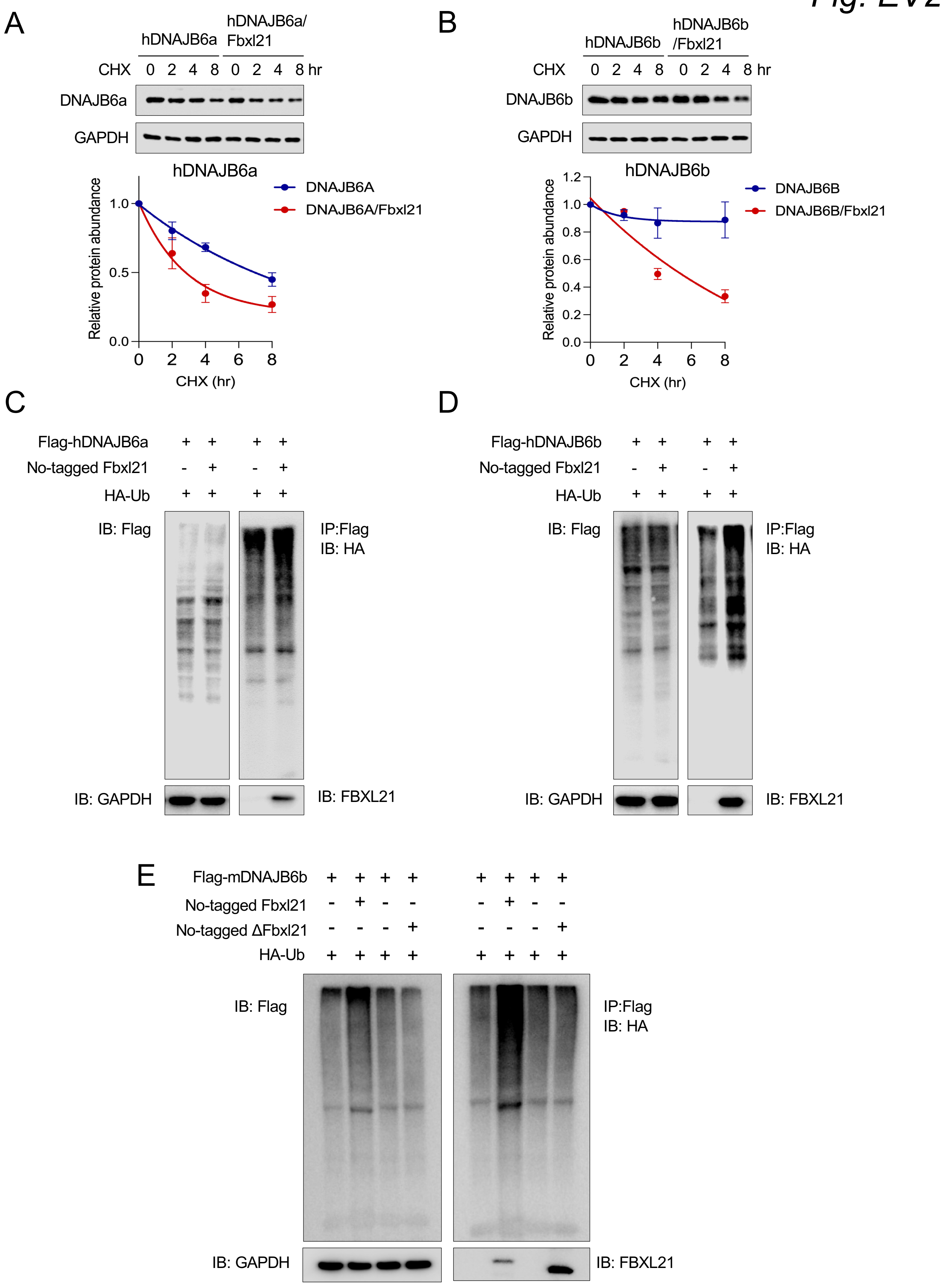
A
hDNAJB6a
hDNAJB6a/ Fbxl21
CHX 0 2 4 8 0 2 4 8 hr
DNAJB6a
GAPDH
hDNAJB6a
DNAJB6A
DNAJB6A/Fbxl21
Relative protein abundance
CHX (hr)
B
hDNAJB6b
hDNAJB6b /Fbxl21
CHX 0 2 4 8 0 2 4 8 hr
DNAJB6b
GAPDH
hDNAJB6b
DNAJB6B
DNAJB6B/Fbxl21
Relative protein abundance
CHX (hr)
C
Flag-hDNAJB6a + + + +
No-tagged Fbxl21 - + - +
HA-Ub + + + +
IB: Flag
IP:Flag
IB: HA
IB: GAPDH
IB: FBXL21
D
Flag-hDNAJB6b + + + +
No-tagged Fbxl21 - + - +
HA-Ub + + + +
IB: Flag
IP:Flag
IB: HA
IB: GAPDH
IB: FBXL21
E
Flag-mDNAJB6b + + + + + + + +
No-tagged Fbxl21 - + - - - + - -
No-tagged ΔFbxl21 - - - + - - - +
HA-Ub + + + + + + + +
IB: Flag
IP:Flag
IB: HA
IB: GAPDH
IB: FBXL21

**Figure EV2. FBXL21 regulates degradation and ubiquitination of human DNAJB6 isoforms.** (**A–B**) Cycloheximide (CHX) chase assays were performed to assess the protein stability of human DNAJB6a (A) and DNAJB6b (**B**) in 293T cells. Protein half-lives were quantified. Half-lives: hDnajb6a, 7.05 h; hDnajb6a/Fbxl21, 2.82 h; hDnajb6b, >24 h; hDnajb6b/Fbxl21. Data are presented as mean ± SEM (n = 3). (**C–D**) FBXL21 promotes the ubiquitination of human DNAJB6 isoforms. Flag-tagged DNAJB6a (**C**) or DNAJB6b (**D**) was co-transfected with untagged FBXL21 and HA-tagged ubiquitin (HA-Ub), followed by ubiquitination assays. (**E**) ΔFBXL21 abolished its ability to ubiquitinate DNAJB6b in 293T cells. Flag-tagged mDNAJB6b was co-transfected with either wild-type FBXL21 or ΔFBXL21 with HA-tagged ubiquitin (HA-Ub), followed by ubiquitination assays. N numbers indicate biological replicates for each sample group, and representative data from three independent experiments are shown.

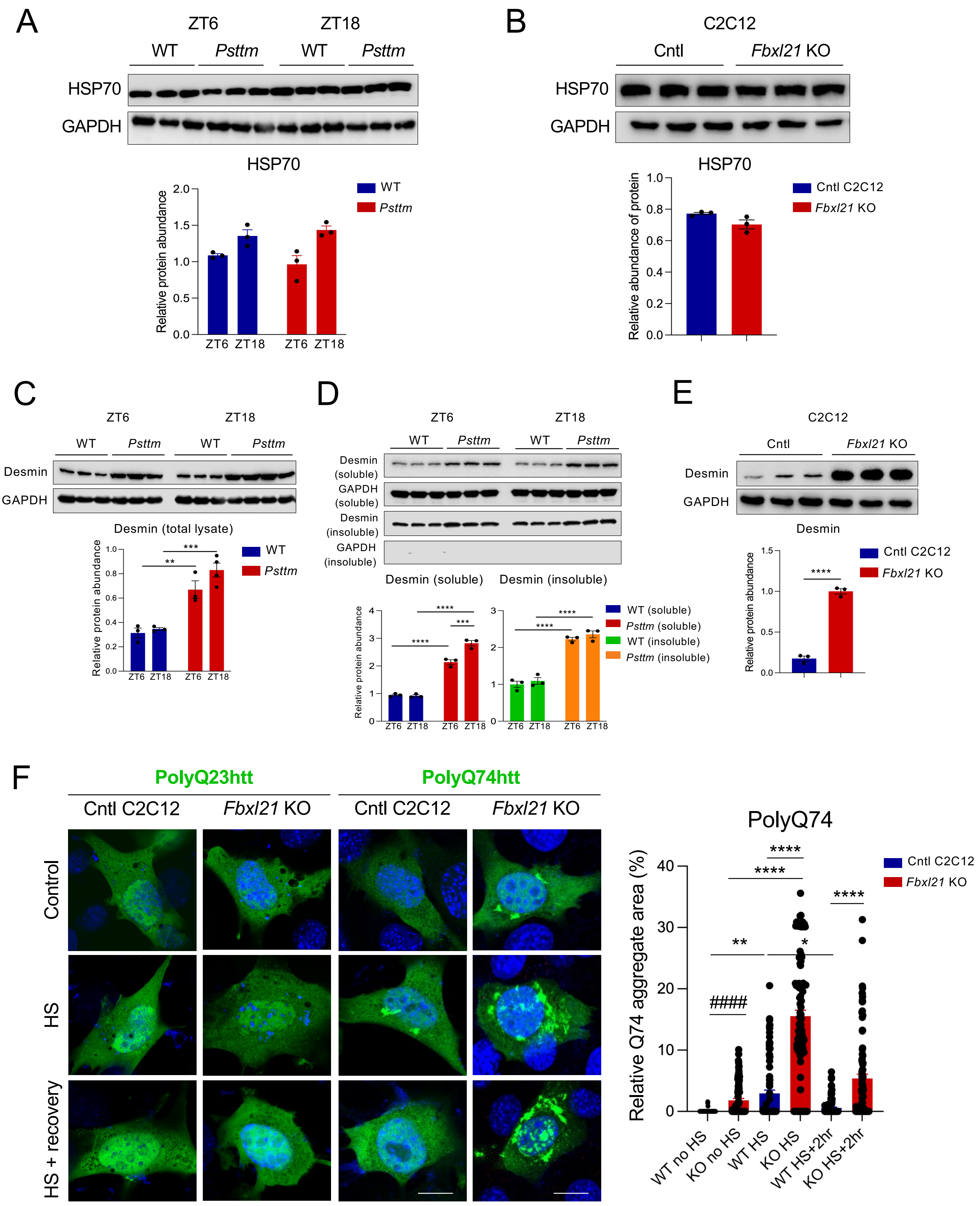
A
ZT6
ZT18
WT
Psttm
WT
Psttm
HSP70
GAPDH
HSP70
Relative protein abundance
2.0
1.5
1.0
0.5
0.0
ZT6 ZT18 ZT6 ZT18
WT
Psttm
B
C2C12
Cntl
Fbxl21 KO
HSP70
GAPDH
HSP70
Relative abundance of protein
1.0
0.8
0.6
0.4
0.2
0.0
Cntl C2C12
Fbxl21 KO
C
ZT6
ZT18
WT
Psttm
WT
Psttm
Desmin
GAPDH
Desmin (total lysate)
Relative protein abundance
1.0
0.8
0.6
0.4
0.2
0.0
***
**
ZT6 ZT18 ZT6 ZT18
WT
Psttm
D
ZT6
ZT18
WT
Psttm
WT
Psttm
Desmin (soluble)
GAPDH (soluble)
Desmin (insoluble)
GAPDH (insoluble)
Desmin (soluble)
Desmin (insoluble)
Relative protein abundance
4
3
2
1
0
****
***
****
ZT6 ZT18 ZT6 ZT18
3
2
1
0
****
****
ZT6 ZT18 ZT6 ZT18
WT (soluble)
Psttm (soluble)
WT (insoluble)
Psttm (insoluble)
E
C2C12
Cntl
Fbxl21 KO
Desmin
GAPDH
Desmin
Relative protein abundance
1.5
1.0
0.5
0.0
****
Cntl C2C12
Fbxl21 KO
F
PolyQ23htt
PolyQ74htt
Cntl C2C12
Fbxl21 KO
Cntl C2C12
Fbxl21 KO
Control
HS
HS + recovery
PolyQ74
Relative Q74 aggregate area (%)
40
30
20
10
0
****
****
****
**
*
####
WT no HS
KO no HS
WT HS
KO HS
WT HS+2hr
KO HS+2hr
Cntl C2C12
Fbxl21 KO

**Figure EV3. FBXL21 regulates desmin proteostasis and polyQ74htt aggregation without altering HSP70 expression.** (**A**) Immunoblot analysis of HSP70 protein levels in wild-type (WT) and *Psttm* mice collected at Zeitgeber time 6 (ZT6; daytime) and ZT18 (nighttime). GAPDH was used as a loading control. Data are presented as mean ± SEM (n = 3/group/time point). No mark indicates not statistically significant. $P$ = 0.7027 (WT ZT6 vs. *Psttm* ZT6) and $P$ = 0.8803 (WT ZT18 vs. *Psttm* ZT18); Two-way ANOVA with Tukey's multiple comparisons test. (**B**) Immunoblot analysis of HSP70 protein levels in control and *Fbxl21* KO C2C1 cells. Data are presented as mean ± SEM (n = 3). No mark indicates not statistically significant. $P$ = 0.0816; Unpaired Student's t-test. (**C**) Immunoblot analysis of Desmin protein levels in WT and *Psttm* mice. GAPDH was used as a loading control. Data are presented as mean ± SEM (n = 3–4). **$P$ = 0.0059 *and* ***$P$ = 0.0004; two-way ANOVA indicates statistical differences between genotypes (WT vs. *Psttm*). (**D**) Immunoblot of Desmin in soluble and insoluble fractions from WT and *Psttm* mice. GAPDH was used as a positive control for the soluble fraction. Data are presented as mean ± SEM (n = 3). ***$P$ = 0.0001 and ****$P$ < 0.0001; Two-way ANOVA with Tukey's multiple comparisons test, comparing between WT and *Psttm* mice or between time points. (**E**) Immunoblot analysis of Desmin protein levels in Cntl and *Fbxl21* KO C2C12 cells. GAPDH was used as a loading control. Data are presented as mean ± SEM (n = 3). ****$P$ < 0.0001; Unpaired Student's t-test. (**F**) Confocal microscopy analysis in Cntl and *Fbxl21* KO C2C12 cells expressing polyQ23htt-EGFP and polyQ74htt-EGFP. PolyQ23htt-EGFP was used as a negative control, and no aggregates were detected in the groups. The relative Q74 aggregate area (%) was quantified using ImageJ. Data are presented as mean ± SEM (n=100 cells/group). *$P$ = 0.0253 (WT HS vs. WT HS recovery), **$P$ = 0.0023 (WT no HS vs. WT HS), and ****$P$ < 0.0001; One-way ANOVA showing significant differences between groups. $^{\#\#\#\#}P$ < 0.0001; Unpaired Student's *t*-test. N numbers indicate biological replicates for each sample group, and representative data from three independent experiments are shown.

### *Fig. EV4*

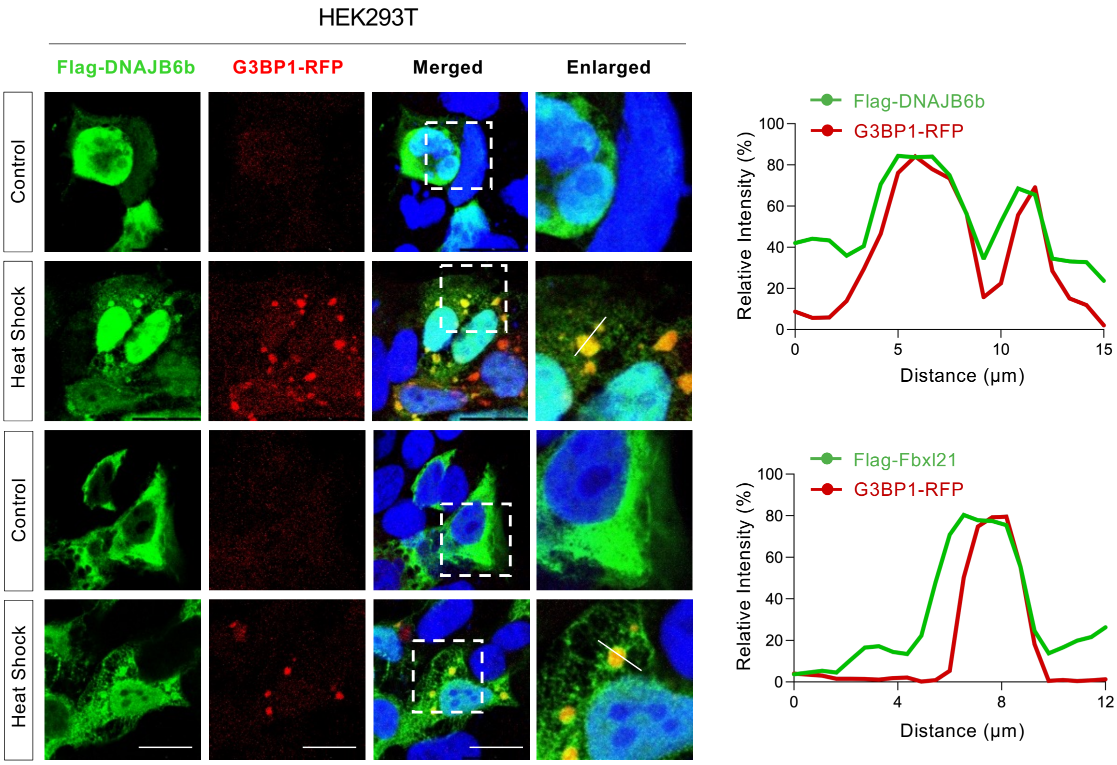

HEK293T
Flag-DNAJB6b
G3BP1-RFP
Merged
Enlarged
Control
Heat Shock
Control
Heat Shock
Flag-DNAJB6b
G3BP1-RFP
Relative Intensity (%)
Distance (μm)
Flag-Fbxl21
G3BP1-RFP
Relative Intensity (%)
Distance (μm)

**Figure EV4. DNAJB6 and FBXL21 colocalize in stress granules. Left panel:** Confocal microscopy of HEK293T cells co-expressing either Flag-DNAJB6b and G3BP1-RFP or Flag-FBXL21 and G3BP1-RFP. Scale bar: 20 µm. **Right panels**: Line profiles showing the mean fluorescence intensities analyzed by using Prism. Representative images are shown from three independent experiments are shown. Scale bar: 20 µm.

### Fig. EV5

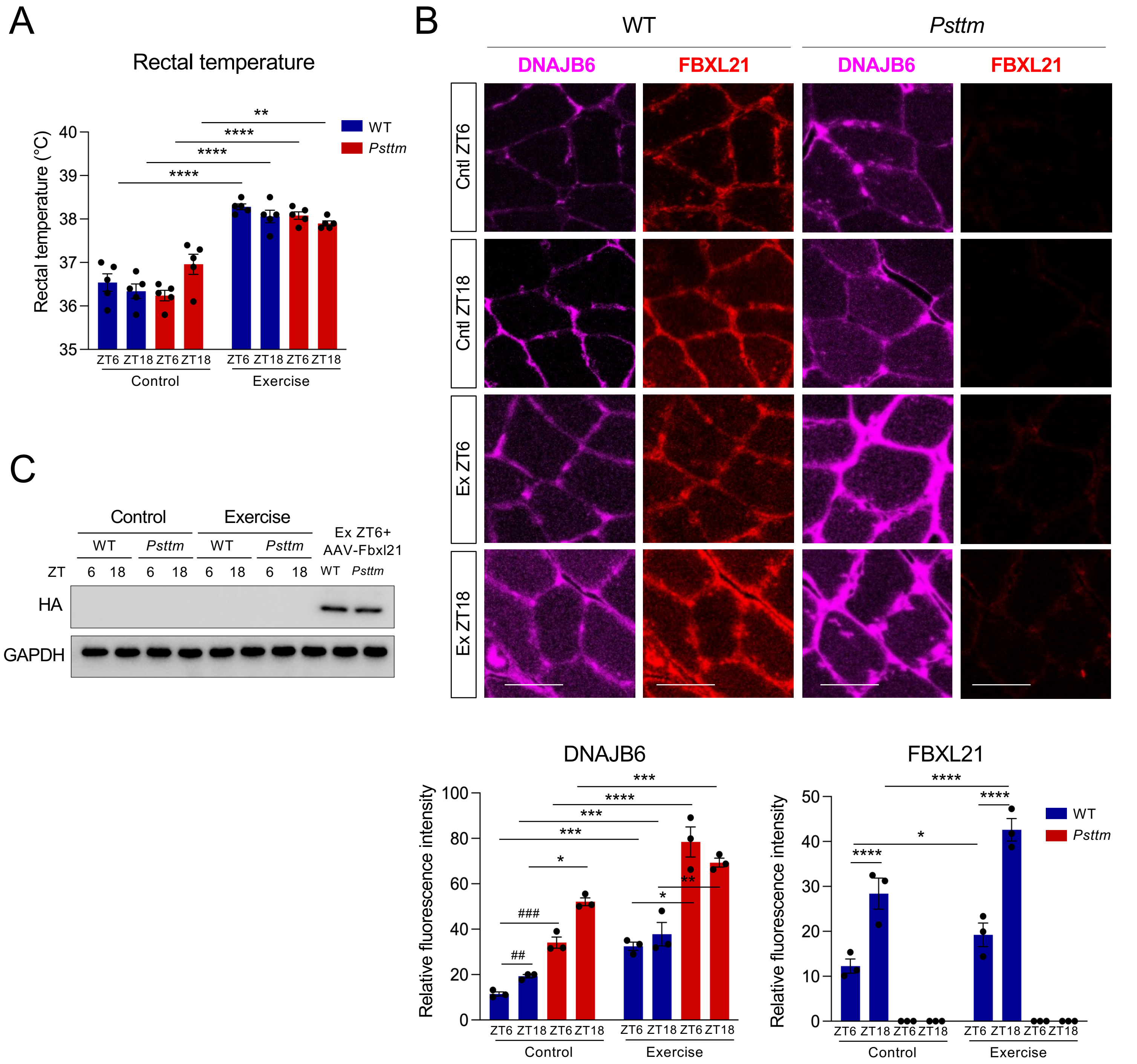
A
Rectal temperature
Rectal temperature (°C)
WT
Psttm
ZT6 ZT18 ZT6 ZT18 ZT6 ZT18 ZT6 ZT18
Control
Exercise
B
WT
Psttm
DNAJB6
FBXL21
Cntl ZT6
Cntl ZT18
Ex ZT6
Ex ZT18
DNAJB6
FBXL21
Relative fluorescence intensity
Control
Exercise
C
Control
Exercise
Ex ZT6+ AAV-Fbxl21
WT Psttm WT Psttm
ZT 6 18 6 18 6 18 6 18 WT Psttm
HA
GAPDH

**Figure EV5. Exercise alters DNAJB6 and FBXL21 expression levels in WT and *Psttm* mice in a time-dependent manner.** (**A**) Rectal temperature measured using a rectal probe (ThermoWorks) at different time points. Data are presented as mean ± SEM (n = 5 mice/group/time point). No mark indicates not statistically significant. $^{**}P = 0.0017$ and $^{****}P < 0.0001$, Two-way ANOVA with Tukey's multiple comparisons test, comparing between groups. (**B**) Immunofluorescence staining of DNAJB6 and FBXL21 in gastrocnemius muscle cross-sections from wild-type (WT) and *Psttm* mice under control (non-exercise) conditions. Scale bar: 20 μm. Lower left panel: Quantification of DNAJB6 fluorescence intensity in gastrocnemius muscle cross-sections from WT and *Psttm* mice under control and acute exercise conditions. Lower right panel: Quantification of FBXL21 fluorescence intensity in gastrocnemius muscle cross-sections from WT and *Psttm* mice under control and acute exercise conditions. Data are presented as mean ± SEM (n = 3/group/time point). For DNAJB6, $^{*}P = 0.0483$ (control WT ZT18 vs. control *Psttm* ZT18), $^{*}P = 0.0131$ (exercise WT ZT6 vs. exercise *Psttm* ZT6), $^{**}P = 0.0061$ (exercise WT ZT18 vs. exercise *Psttm* ZT18), $^{***}P = 0.0015$ (control WT ZT6 vs. exercise WT ZT6), $^{***}P = 0.0001$ (exercise WT ZT18 vs. exercise *Psttm* ZT18), and $^{****}P < 0.0001$; Two-way ANOVA with Tukey's multiple comparisons test. $^{\#\#}P = 0.0028$ and $^{\#\#\#}P = 0.0010$; Unpaired Student's *t*-test. For FBXL21, $^{*}P = 0.0176$ and $^{****}P < 0.0001$; Two-way ANOVA with Tukey's multiple comparisons test. (**C**) Immunoblot assay for HA detection in gastrocnemius tissues of WT and *Psttm* mice subjected to daytime exercise and AAV-CK6-Fbxl21-HA injection.

**Table EV1.** HEK293T cells were transfected with a control or Flag-FBXL21 vector. Immunoprecipitation was performed by using anti-Flag beads, followed by mass spectrometry. The list of proteins identified is shown.

| GeneID | GeneSymbol |
|---|---|
| 10049 | DNAJB6 |
| 1408 | CRY2 |
| 5692 | PSMB4 |
| 7917 | BAG6 |
| 5689 | PSMB1 |
| 5688 | PSMA7 |
| 1407 | CRY1 |
| 5690 | PSMB2 |
| 5714 | PSMD8 |
| 5694 | PSMB6 |
| 8878 | SQSTM1 |
| 5693 | PSMB5 |
| 8079 | MLF2 |
| 9532 | BAG2 |
| 11080 | DNAJB4 |
| 9500 | MAGED1 |
| 23552 | CDK20 |
| 2268 | FGR |
| 7525 | YES1 |
| 2534 | FYN |
| 640 | BLK |
| 3932 | LCK |
| 3315 | HSPB1 |
| 3055 | HCK |
| 4067 | LYN |
| 4117 | MAK |
| 10910 | SUGT1 |
| 2264 | FGFR4 |
| 2263 | FGFR2 |
| 2260 | FGFR1 |
| 22858 | ICK |
| 5700 | PSMC1 |
| 2261 | FGFR3 |
| 8471 | IRS4 |
| 10916 | MAGED2 |
| 5701 | PSMC2 |
| 5686 | PSMA5 |
| 10294 | DNAJA2 |

| | |
|---|---|
| 3875 | KRT18 |
| 5710 | PSMD4 |
| 5717 | PSMD11 |
| 4741 | NEFM |
| 3310 | HSPA6 |
| 9967 | THRAP3 |
| 6500 | SKP1 |
| 5708 | PSMD2 |
| 1719 | DHFR |
| 5707 | PSMD1 |
| 5706 | PSMC6 |
| 5682 | PSMA1 |
| 9861 | PSMD6 |
| 5704 | PSMC4 |
| 63929 | XPNPEP3 |
| 5705 | PSMC5 |
| 3301 | DNAJA1 |
| 346288 | 45914 |
| 10213 | PSMD14 |
| 5713 | PSMD7 |
| 5930 | RBBP6 |
| 5709 | PSMD3 |
| 23176 | 45908 |
| 23157 | 45906 |
| 5702 | PSMC3 |
| 10657 | KHDRBS1 |
| 100529097 | RPL36A-HNRNPH2 |
| 6173 | RPL36A |
| 6229 | RPS24 |
| 10963 | STIP1 |
| 7415 | VCP |
| 4001 | LMNB1 |
| 6949 | TCOF1 |
| 6921 | TCEB1 |
| 6135 | RPL11 |
| 989 | 45907 |
| 8668 | EIF3I |
| 55752 | 45911 |
| 5591 | PRKDC |
| 9045 | RPL14 |
| 10574 | CCT7 |
| 9131 | AIFM1 |
| 10197 | PSME3 |
| 6189 | RPS3A |

| | |
|---|---|
| 8531 | YBX3 |
| 23020 | SNRNP200 |
| 6136 | RPL12 |
| 3182 | HNRNPAB |
| 3608 | ILF2 |
| 22948 | CCT5 |
| 8407 | TAGLN2 |
| 3837 | KPNB1 |
| 3308 | HSPA4 |
| 1213 | CLTC |
| 716 | C1S |
| 5648 | MASP1 |
| 810 | CALML3 |
| 2784 | GNB3 |
| 3857 | KRT9 |
| 4831 | NME2 |
| 3849 | KRT2 |
| 3848 | KRT1 |
| 3858 | KRT10 |
| 10929 | SRSF8 |
| 100996747 | LOC100996747 |
| 6152 | RPL24 |
| 10381 | TUBB3 |
| 3854 | KRT6B |
| 5685 | PSMA4 |
| 8266 | UBL4A |
| 143471 | PSMA8 |
| 6637 | SNRPG |
| 3107 | HLA-C |
| 5691 | PSMB3 |
| 3106 | HLA-B |
| 5687 | PSMA6 |
| 51608 | GET4 |
| 5684 | PSMA3 |
| 7699 | ZNF140 |
| 3337 | DNAJB1 |
| 5683 | PSMA2 |
| 140801 | RPL10L |
| 143244 | EIF5AL1 |
| 286887 | KRT6C |
| 3135 | HLA-G |
| 10456 | HAX1 |
| 51377 | UCHL5 |
| 714 | C1QC |

| | |
|---|---|
| 5437 | POLR2H |
| 196374 | KRT78 |
| 6749 | SSRP1 |
| 123283 | TARSL2 |
| 1019 | CDK4 |
| 372 | ARCN1 |
| 8087 | FXR1 |
| 10273 | STUB1 |
| 2332 | FMR1 |
| 3853 | KRT6A |
| 1025 | CDK9 |
| 1160 | CKMT2 |
| 9513 | FXR2 |
| 10075 | HUWE1 |
| 50628 | GEMIN4 |
| 7157 | TP53 |
| 1832 | DSP |
| 3851 | KRT4 |
| 2949 | GSTM5 |
| 3868 | KRT16 |
| 2946 | GSTM2 |
| 3852 | KRT5 |
| 8340 | HIST1H2BL |
| 2335 | FN1 |
| 3188 | HNRNPH2 |
| 200895 | DHFRL1 |
| 6233 | RPS27A |
| 7311 | UBA52 |
| 7314 | UBB |
| 7316 | UBC |
| 5837 | PYGM |
| 9919 | SEC16A |
| 9991 | PTBP3 |
| 10643 | IGF2BP3 |
| 983 | CDK1 |
| 388697 | HRNR |
| 5042 | PABPC3 |
| 10383 | TUBB4B |
| 56648 | EIF5A2 |
| 5573 | PRKAR1A |
| 5224 | PGAM2 |
| 55082 | ARGLU1 |
| 2483 | FRG1 |
| 102288414 | C11orf98 |

| | |
|---|---|
| 9584 | RBM39 |
| 3312 | HSPA8 |
| 3043 | HBB |
| 3045 | HBD |
| 3861 | KRT14 |
| 5718 | PSMD12 |
| 480 | ATP1A4 |
| 7453 | WARS |
| 7266 | DNAJC7 |
| 6170 | RPL39 |
| 8148 | TAF15 |
| 478 | ATP1A3 |
| 477 | ATP1A2 |
| 11047 | ADRM1 |
| 495 | ATP4A |
| 479 | ATP12A |
| 3047 | HBG1 |
| 3048 | HBG2 |
| 3046 | HBE1 |
| 3303 | HSPA1A |
| 3304 | HSPA1B |
| 3008 | HIST1H1E |
| 3007 | HIST1H1D |
| 6235 | RPS29 |
| 1073 | CFL2 |
| 79169 | C1orf35 |
| 388524 | RPSAP58 |
| 4841 | NONO |
| 1654 | DDX3X |
| 9470 | EIF4E2 |
| 8721 | EDF1 |
| 200916 | RPL22L1 |
| 84617 | TUBB6 |
| 790 | CAD |
| 718 | C3 |
| 682 | BSG |
| 2197 | FAU |
| 6635 | SNRPE |
| 128312 | HIST3H2BB |
| 5695 | PSMB7 |
| 51639 | SF3B6 |
| 79576 | NKAP |
| 3039 | HBA1 |
| 3040 | HBA2 |

| | |
|---|---|
| 3856 | KRT8 |
| 3860 | KRT13 |
| 25873 | RPL36 |
| 2 | A2M |
| 1029 | CDKN2A |
| 5962 | RDX |
| 646817 | SETSIP |
| 117159 | DCD |
| 6218 | RPS17 |
| 292 | SLC25A5 |
| 2244 | FGB |
| 55740 | ENAH |
| 100423062 | IGLL5 |
| 388698 | FLG2 |
| 6421 | SFPQ |
| 6232 | RPS27 |
| 51121 | RPL26L1 |
| 94239 | H2AFV |
| 3015 | H2AFZ |
| 6228 | RPS23 |
| 51428 | DDX41 |
| 6142 | RPL18A |
| 9349 | RPL23 |
| 4670 | HNRNPM |
| 6168 | RPL37A |
| 6207 | RPS13 |
| 440093 | H3F3C |
| 440686 | HIST2H3PS2 |
| 6748 | SSR4 |
| 6138 | RPL15 |
| 1665 | DHX15 |
| 6280 | S100A9 |
| 121504 | HIST4H4 |
| 554313 | HIST2H4B |
| 8294 | HIST1H4I |
| 8359 | HIST1H4A |
| 8360 | HIST1H4D |
| 8361 | HIST1H4F |
| 8362 | HIST1H4K |
| 8363 | HIST1H4J |
| 8364 | HIST1H4C |
| 8365 | HIST1H4H |
| 8366 | HIST1H4B |
| 8367 | HIST1H4E |

| | |
|---|---|
| 8368 | HIST1H4L |
| 8370 | HIST2H4A |
| 2266 | FGG |
| 7431 | VIM |
| 26058 | GIGYF2 |
| 3018 | HIST1H2BB |
| 8348 | HIST1H2BO |
| 8349 | HIST2H2BE |
| 8970 | HIST1H2BJ |
| 6157 | RPL27A |
| 335 | APOA1 |
| 5719 | PSMD13 |
| 3014 | H2AFX |
| 3320 | HSP90AA1 |
| 387082 | SUMO4 |
| 6612 | SUMO3 |
| 5861 | RAB1A |
| 6634 | SNRPD3 |
| 6241 | RRM2 |
| 203068 | TUBB |
| 6122 | RPL3 |
| 6222 | RPS18 |
| 6144 | RPL21 |
| 6169 | RPL38 |
| 5110 | PCMT1 |
| 3698 | ITIH2 |
| 6165 | RPL35A |
| 6124 | RPL4 |
| 80086 | TUBA4B |
| 10131 | TRAP1 |
| 7846 | TUBA1A |
| 6194 | RPS6 |
| 6141 | RPL18 |
| 3326 | HSP90AB1 |
| 213 | ALB |
| 3313 | HSPA9 |
| 6224 | RPS20 |
| 55506 | H2AFY2 |
| 3006 | HIST1H1C |
| 6160 | RPL31 |
| 29767 | TMOD2 |
| 720 | C4A |
| 100293534 | C4B_2 |
| 110384692 | LOC110384692 |

| | |
|---|---|
| 721 | C4B |
| 5887 | RAD23B |
| 6230 | RPS25 |
| 6130 | RPL7A |
| 6208 | RPS14 |
| 3309 | HSPA5 |
| 6203 | RPS9 |
| 9987 | HNRNPDL |
| 6143 | RPL19 |
| 3295 | HSD17B4 |
| 509 | ATP5C1 |
| 6137 | RPL13 |
| 126961 | HIST2H3C |
| 3020 | H3F3A |
| 3021 | H3F3B |
| 333932 | HIST2H3A |
| 653604 | HIST2H3D |
| 8290 | HIST3H3 |
| 8350 | HIST1H3A |
| 8351 | HIST1H3D |
| 8352 | HIST1H3C |
| 8353 | HIST1H3E |
| 8354 | HIST1H3I |
| 8355 | HIST1H3G |
| 8356 | HIST1H3J |
| 8357 | HIST1H3H |
| 8358 | HIST1H3B |
| 8968 | HIST1H3F |
| 6164 | RPL34 |
| 2079 | ERH |
| 6187 | RPS2 |
| 6159 | RPL29 |
| 140735 | DYNLL2 |
| 8655 | DYNLL1 |
| 11224 | RPL35 |
| 1267 | CNP |
| 6205 | RPS11 |
| 6206 | RPS12 |
| 10911 | UTS2 |
| 6217 | RPS16 |
| 6202 | RPS8 |
| 10376 | TUBA1B |
| 84790 | TUBA1C |
| 23521 | RPL13A |

| | |
|---|---|
| 6210 | RPS15A |
| 23119 | HIC2 |
| 5831 | PYCR1 |
| 54734 | RAB39A |
| 6191 | RPS4X |
| 4736 | RPL10A |
| 6188 | RPS3 |
| 55127 | HEATR1 |
| 1460 | CSNK2B |
| 3017 | HIST1H2BD |
| 440689 | HIST2H2BF |
| 102724334 | LOC102724334 |
| 54145 | H2BFS |
| 8339 | HIST1H2BG |
| 8341 | HIST1H2BN |
| 8342 | HIST1H2BM |
| 8343 | HIST1H2BF |
| 8344 | HIST1H2BE |
| 8345 | HIST1H2BH |
| 8346 | HIST1H2BI |
| 8347 | HIST1H2BC |
| 85236 | HIST1H2BK |
| 102724594 | U2AF1L5 |
| 7307 | U2AF1 |
| 1973 | EIF4A1 |
| 3005 | H1F0 |
| 6132 | RPL8 |
| 6129 | RPL7 |
| 3066 | HDAC2 |
| 3065 | HDAC1 |
| 4809 | SNU13 |
| 1915 | EEF1A1 |
| 5931 | RBBP7 |
| 6166 | RPL36AL |
| 6155 | RPL27 |
| 6201 | RPS7 |
| 908 | CCT6A |
| 6133 | RPL9 |
| 9774 | BCLAF1 |
| 6193 | RPS5 |
| 26986 | PABPC1 |
| 6209 | RPS15 |
| 4173 | MCM4 |
| 3178 | HNRNPA1 |

| | |
|---|---|
| 4926 | NUMA1 |
| 6223 | RPS19 |
| 9343 | EFTUD2 |
| 285672 | SREK1IP1 |
| 2243 | FGA |
| 3191 | HNRNPL |
| 6146 | RPL22 |
| 6181 | RPLP2 |
| 6147 | RPL23A |
| 6154 | RPL26 |
| 23481 | PES1 |
| 6613 | SUMO2 |
| 10575 | CCT4 |
| 30000 | TNPO2 |
| 55379 | LRRC59 |
| 83452 | RAB33B |
| 5928 | RBBP4 |
| 708 | C1QBP |
| 10694 | CCT8 |
| 2664 | GDI1 |
| 4673 | NAP1L1 |
| 51441 | YTHDF2 |
| 84823 | LMNB2 |
| 3181 | HNRNPA2B1 |
| 3376 | IARS |
| 60 | ACTB |
| 71 | ACTG1 |
| 1964 | EIF1AX |
| 9086 | EIF1AY |
| 10576 | CCT2 |
| 3183 | HNRNPC |
| 1655 | DDX5 |
| 6175 | RPLP0 |
| 4869 | NPM1 |
| 58 | ACTA1 |
| 70 | ACTC1 |
| 3838 | KPNA2 |
| 2782 | GNB1 |
| 2783 | GNB2 |
| 59345 | GNB4 |
| 4282 | MIF |
| 6119 | RPA3 |
| 51560 | RAB6B |
| 5870 | RAB6A |

| | |
|---|---|
| 4116 | MAGOH |
| 55110 | MAGOHB |
| 7203 | CCT3 |
| 4624 | MYH6 |
| 9782 | MATR3 |
| 4625 | MYH7 |
| 100996928 | C7orf55-LUC7L2 |
| 6158 | RPL28 |
| 3192 | HNRNPU |
| 29796 | UQCR10 |
| 23215 | PRRC2C |
| 1933 | EEF1B2 |
| 6240 | RRM1 |
| 6950 | TCP1 |
| 5902 | RANBP1 |
| 498 | ATP5A1 |
| 8607 | RUVBL1 |
| 10134 | BCAP31 |
| 6231 | RPS26 |
| 1937 | EEF1G |
| 8450 | CUL4B |
| 5478 | PPIA |
| 5725 | PTBP1 |
| 90850 | ZNF598 |
| 23170 | TTLL12 |
| 3945 | LDHB |
| 6161 | RPL32 |
| 142 | PARP1 |
| 10594 | PRPF8 |
| 5901 | RAN |
| 3329 | HSPD1 |
| 9793 | CKAP5 |
| 10419 | PRMT5 |
| 808 | CALM3 |
| 801 | CALM1 |
| 805 | CALM2 |
| 1537 | CYC1 |
| 27316 | RBMX |
| 1938 | EEF2 |
| 4629 | MYH11 |
| 23256 | SCFD1 |
| 6125 | RPL5 |
| 3069 | HDLBP |
| 5052 | PRDX1 |

| | |
|---|---|
| 3187 | HNRNPH1 |
| 10642 | IGF2BP1 |
| 6128 | RPL6 |
| 10399 | GNB2L1 |
| 10606 | PAICS |
| 2739 | GLO1 |
| 7295 | TXN |
| 5631 | PRPS1 |
| 1639 | DCTN1 |
| 2547 | XRCC6 |
| 4666 | NACA |
| 10521 | DDX17 |
| 220988 | HNRNPA3 |
| 376267 | RAB15 |
| 3190 | HNRNPK |
| 5093 | PCBP1 |
| 7094 | TLN1 |
| 2023 | ENO1 |
| 2597 | GAPDH |
| 515 | ATP5F1 |
| 3939 | LDHA |
| 26135 | SERBP1 |
| 7534 | YWHAZ |
| 6059 | ABCE1 |
| 506 | ATP5B |
| 6902 | TBCA |
| 5216 | PFN1 |
| 10856 | RUVBL2 |
| 5230 | PGK1 |
| 11331 | PHB2 |
| 6139 | RPL17 |
| 100526842 | RPL17-C18orf32 |
| 10189 | ALYREF |
| 10155 | TRIM28 |
| 4513 | COX2 |
| 7419 | VDAC3 |
| 800 | CALD1 |
| 1786 | DNMT1 |
| 5905 | RANGAP1 |
| 5315 | PKM |
| 5250 | SLC25A3 |
| 5161 | PDHA2 |
| 7416 | VDAC1 |
| 28988 | DBNL |

| | |
|---|---|
| 4691 | NCL |
| 100529239 | RPS10-NUDT3 |
| 7167 | TPI1 |
| 55621 | TRMT1 |
| 9588 | PRDX6 |
| 1152 | CKB |
| 9040 | UBE2M |
| 9554 | SEC22B |
| 7531 | YWHAE |
| 5929 | RBBP5 |
| 5886 | RAD23A |
| 9184 | BUB3 |
| 191 | AHCY |
| 64062 | RBM26 |
| 378 | ARF4 |
| 23223 | RRP12 |
| 5479 | PPIB |
| 226 | ALDOA |
| 1968 | EIF2S3 |
| 230 | ALDOC |
| 255308 | LOC255308 |
| 93974 | ATPIF1 |
| 9590 | AKAP12 |
| 9555 | H2AFY |
| 6134 | RPL10 |
| 2618 | GART |
| 6184 | RPN1 |
| 1159 | CKMT1B |
| 6520 | SLC3A2 |
| 548596 | CKMT1A |
| 6730 | SRP68 |
| 6638 | SNRPN |
| 6628 | SNRPB |
| 65108 | MARCKSL1 |
| 11140 | CDC37 |
| 55037 | PTCD3 |
| 55749 | CCAR1 |
| 3796 | KIF2A |
| 10988 | METAP2 |
| 27101 | CACYBP |
| 6510 | SLC1A5 |
| 22938 | SNW1 |
| 8801 | SUCLG2 |
| 440 | ASNS |

| | |
|---|---|
| 2058 | EPRS |
| 55920 | RCC2 |
| 6204 | RPS10 |
| 3146 | HMGB1 |
| 4144 | MAT2A |
| 4904 | YBX1 |
| 51631 | LUC7L2 |
| 6897 | TARS |
| 7430 | EZR |
| 4637 | MYL6 |
| 26227 | PHGDH |
| 7170 | TPM3 |
| 9126 | SMC3 |
| 2521 | FUS |
| 54039 | PCBP3 |
| 1981 | EIF4G1 |
| 3842 | TNPO1 |
| 10726 | NUDC |
| 3251 | HPRT1 |
| 821 | CANX |
| 5223 | PGAM1 |
| 22803 | XRN2 |
| 11338 | U2AF2 |
| 5034 | P4HB |
| 760 | CA2 |
| 80184 | CEP290 |
| 1965 | EIF2S1 |
| 7532 | YWHAG |
| 1936 | EEF1D |
| 56829 | ZC3HAV1 |
| 27044 | SND1 |
| 2317 | FLNB |
| 5868 | RAB5A |
| 5499 | PPP1CA |
| 7417 | VDAC2 |
| 6767 | ST13 |
| 6051 | RNPEP |
| 5859 | QARS |
| 654364 | NME1-NME2 |
| 988 | CDC5L |
| 117246 | FTSJ3 |
| 302 | ANXA2 |
| 9879 | DDX46 |
| 5869 | RAB5B |

| | |
|---|---|
| 92140 | MTDH |
| 8653 | DDX3Y |
| 55705 | IPO9 |
| 1660 | DHX9 |
| 2810 | SFN |
| 4478 | MSN |
| 11252 | PACSIN2 |
| 4000 | LMNA |
| 2821 | GPI |
| 3921 | RPSA |
| 26121 | PRPF31 |
| 54205 | CYCS |
| 10935 | PRDX3 |
| 5757 | PTMA |
| 7965 | AIMP2 |
| 7879 | RAB7A |
| 9218 | VAPA |
| 6711 | SPTBN1 |
| 50809 | HP1BP3 |
| 81570 | CLPB |
| 4830 | NME1 |
| 1278 | COL1A2 |
| 7247 | TSN |
| 54815 | GATAD2A |
| 3336 | HSPE1 |
| 79784 | MYH14 |
| 4860 | PNP |
| 9092 | SART1 |
| 6238 | RRBP1 |
| 6427 | SRSF2 |
| 1642 | DDB1 |
| 2665 | GDI2 |
| 7284 | TUFM |
| 163 | AP2B1 |
| 5245 | PHB |
| 57459 | GATAD2B |
| 4522 | MTHFD1 |
| 204 | AK2 |
| 2584 | GALK1 |
| 56945 | MRPS22 |
| 55738 | ARFGAP1 |
| 8604 | SLC25A12 |
| 4175 | MCM6 |
| 10989 | IMMT |

| | |
|---|---|
| 2947 | GSTM3 |
| 27339 | PRPF19 |
| 7112 | TMPO |
| 523 | ATP6V1A |
| 10514 | MYBBP1A |
| 4628 | MYH10 |
| 7001 | PRDX2 |
| 6709 | SPTAN1 |
| 4191 | MDH2 |
| 3145 | HMBS |
| 7520 | XRCC5 |
| 10130 | PDIA6 |
| 7175 | TPR |
| 6729 | SRP54 |
| 1653 | DDX1 |
| 10165 | SLC25A13 |
| 276 | AMY1A |
| 277 | AMY1B |
| 278 | AMY1C |
| 279 | AMY2A |
| 280 | AMY2B |
| 10528 | NOP56 |
| 5836 | PYGL |
| 6123 | RPL3L |
| 9114 | ATP6V0D1 |
| 3927 | LASP1 |
| 102724560 | CBSL |
| 814 | CAMK4 |
| 164 | AP1G1 |
| 3416 | IDE |
| 1982 | EIF4G2 |
| 10072 | DPP3 |
| 9097 | USP14 |
| 1632 | ECI1 |
| 47 | ACLY |
| 875 | CBS |
| 5536 | PPP5C |
| 10250 | SRRM1 |
| 3930 | LBR |
| 9688 | NUP93 |
| 100529063 | BCL2L2-PABPN1 |
| 10949 | HNRNPA0 |
| 84991 | RBM17 |
| 23450 | SF3B3 |

| | |
|---|---|
| 84717 | HDGFRP2 |
| 1615 | DARS |
| 373156 | GSTK1 |
| 57510 | XPO5 |
| 1984 | EIF5A |
| 27336 | HTATSF1 |
| 6050 | RNH1 |
| 50814 | NSDHL |
| 7184 | HSP90B1 |
| 8894 | EIF2S2 |
| 5936 | RBM4 |
| 9775 | EIF4A3 |
| 2956 | MSH6 |
| 55004 | LAMTOR1 |
| 10970 | CKAP4 |
| 1315 | COPB1 |
| 51503 | CWC15 |
| 7332 | UBE2L3 |
| 5214 | PFKP |
| 56902 | PNO1 |
| 83540 | NUF2 |
| 2194 | FASN |
| 8570 | KHSRP |
| 55646 | LYAR |
| 9601 | PDIA4 |
| 10801 | 45909 |
| 4176 | MCM7 |
| 2673 | GFPT1 |
| 55660 | PRPF40A |
| 7037 | TFRC |
| 79624 | ARMT1 |
| 65260 | COA7 |
| 1277 | COL1A1 |
| 1072 | CFL1 |
| 5636 | PRPSAP2 |
| 5036 | PA2G4 |
| 874 | CBR3 |
| 23524 | SRRM2 |
| 7812 | CSDE1 |
| 4677 | NARS |
| 10061 | ABCF2 |
| 10969 | EBNA1BP2 |
| 5635 | PRPSAP1 |
| 11198 | SUPT16H |

| | |
|---|---|
| 4436 | MSH2 |
| 8106 | PABPN1 |
| 55840 | EAF2 |
| 100526737 | RBM14-RBM4 |
| 39 | ACAT2 |
| 2288 | FKBP4 |
| 2631 | GBAS |
| 4043 | LRPAP1 |
| 28969 | BZW2 |
| 2319 | FLOT2 |
| 3954 | LETM1 |
| 55347 | ABHD10 |
| 10980 | COPS6 |
| 10212 | DDX39A |
| 51593 | SRRT |
| 197259 | MLKL |
| 10592 | SMC2 |
| 7411 | VBP1 |
| 54499 | TMCO1 |
| 10985 | GCN1 |
| 5226 | PGD |
| 23435 | TARDBP |
| 79869 | CPSF7 |
| 2969 | GTF2I |
| 57136 | APMAP |
| 223 | ALDH9A1 |
| 6917 | TCEA1 |
| 3609 | ILF3 |
| 56993 | TOMM22 |
| 5879 | RAC1 |
| 128240 | APOA1BP |
| 396 | ARHGDIA |
| 6418 | SET |
| 51260 | PBDC1 |
| 4218 | RAB8A |
| 4925 | NUCB2 |
| 6449 | SGTA |
| 5501 | PPP1CC |
| 55119 | PRPF38B |
| 100528064 | NEDD8-MDP1 |
| 51635 | DHRS7 |
| 5160 | PDHA1 |
| 5500 | PPP1CB |
| 23708 | GSPT2 |

| | |
|---|---|
| 11269 | DDX19B |
| 387522 | TMEM189-UBE2V1 |
| 310 | ANXA7 |
| 6624 | FSCN1 |
| 64841 | GNPNAT1 |
| 100037417 | DDTL |
| 22894 | DIS3 |
| 7414 | VCL |
| 1743 | DLST |
| 8971 | H1FX |
| 2935 | GSPT1 |
| 5880 | RAC2 |
| 5881 | RAC3 |
| 4190 | MDH1 |
| 57470 | LRRC47 |
| 301 | ANXA1 |
| 1778 | DYNC1H1 |
| 55176 | SEC61A2 |
| 10890 | RAB10 |
| 55308 | DDX19A |
| 10284 | SAP18 |
| 1652 | DDT |
| 65980 | BRD9 |
| 4171 | MCM2 |
| 735 | C9 |
| 2923 | PDIA3 |
| 80777 | CYB5B |
| 23479 | ISCU |
| 11332 | ACOT7 |
| 23760 | PITPNB |
| 1841 | DTYMK |
| 6281 | S100A10 |
| 10920 | COPS8 |
| 23382 | AHCYL2 |
| 10549 | PRDX4 |
| 2017 | CTTN |
| 10768 | AHCYL1 |
| 58517 | RBM25 |
| 128866 | CHMP4B |
| 5872 | RAB13 |
| 55737 | VPS35 |
| 10452 | TOMM40 |
| 6642 | SNX1 |
| 51762 | RAB8B |

| | |
|---|---|
| 9188 | DDX21 |
| 4172 | MCM3 |
| 1207 | CLNS1A |
| 11100 | HNRNPUL1 |
| 6643 | SNX2 |
| 22824 | HSPA4L |
| 1656 | DDX6 |
| 5211 | PFKL |
| 29920 | PYCR2 |
| 2873 | GPS1 |
| 10121 | ACTR1A |
| 9375 | TM9SF2 |
| 3098 | HK1 |
| 80155 | NAA15 |
| 9130 | FAM50A |
| 54888 | NSUN2 |
| 2052 | EPHX1 |
| 23520 | ANP32C |
| 9898 | UBAP2L |
| 4141 | MARS |
| 9689 | BZW1 |
| 51042 | ZNF593 |
| 87 | ACTN1 |
| 6470 | SHMT1 |
| 2617 | GARS |
| 10007 | GNPDA1 |
| 55157 | DARS2 |
| 51504 | TRMT112 |
| 22820 | COPG1 |
| 10469 | TIMM44 |
| 10120 | ACTR1B |
| 23519 | ANP32D |
| 102723407 | LOC102723407 |
| 3032 | HADHB |
| 8939 | FUBP3 |
| 4085 | MAD2L1 |
| 5236 | PGM1 |
| 5198 | PFAS |
| 51727 | CMPK1 |
| 10682 | EBP |
| 10525 | HYOU1 |
| 1974 | EIF4A2 |
| 50 | ACO2 |
| 51495 | HACD3 |

| | |
|---|---|
| 27068 | PPA2 |
| 26156 | RSL1D1 |
| 1676 | DFFA |
| 5720 | PSME1 |
| 1382 | CRABP2 |
| 3030 | HADHA |
| 9524 | TECR |
| 80218 | NAA50 |
| 2805 | GOT1 |
| 57103 | TIGAR |
| 5747 | PTK2 |
| 5471 | PPAT |
| 80021 | TMEM62 |
| 1983 | EIF5 |
| 90411 | MCFD2 |
| 826 | CAPNS1 |
| 9276 | COPB2 |
| 1314 | COPA |
| 2029 | ENSA |
| 8260 | NAA10 |
| 3093 | UBE2K |
| 55968 | NSFL1C |
| 6429 | SRSF4 |
| 10946 | SF3A3 |
| 476 | ATP1A1 |
| 9295 | SRSF11 |
| 7511 | XPNPEP1 |
| 400 | ARL1 |
| 4733 | DRG1 |
| 8209 | C21orf33 |
| 26355 | FAM162A |
| 8669 | EIF3J |
| 1503 | CTPS1 |
| 102724023 | LOC102724023 |
| 7335 | UBE2V1 |
| 10055 | SAE1 |
| 2764 | GMFB |
| 9669 | EIF5B |
| 9318 | COPS2 |
| 7317 | UBA1 |
| 4076 | CAPRIN1 |
| 115207 | KCTD12 |
| 51520 | LARS |
| 59285 | CACNG6 |

| | |
|---|---|
| 2316 | FLNA |
| 9535 | GMFG |
| 10128 | LRPPRC |
| 4905 | NSF |
| 30001 | ERO1A |
| 10987 | COPS5 |
| 552900 | BOLA2 |
| 8402 | SLC25A11 |
| 8520 | HAT1 |
| 501 | ALDH7A1 |
| 2027 | ENO3 |
| 6342 | SCP2 |
| 7407 | VARS |
| 51501 | C11orf73 |
| 51747 | LUC7L3 |
| 7913 | DEK |
| 55847 | CISD1 |
| 10808 | HSPH1 |
| 10056 | FARSB |
| 494115 | RBMXL1 |
| 125963 | OR1M1 |
| 2885 | GRB2 |
| 94081 | SFXN1 |
| 9550 | ATP6V1G1 |
| 1478 | CSTF2 |
| 10313 | RTN3 |
| 10291 | SF3A1 |
| 3843 | IPO5 |
| 7384 | UQCRC1 |
| 2787 | GNG5 |
| 2107 | ETF1 |
| 81892 | SLIRP |
| 11164 | NUDT5 |
| 1155 | TBCB |
| 158 | ADSL |
| 55970 | GNG12 |
| 4738 | NEDD8 |
| 8663 | EIF3C |
| 51031 | GLOD4 |
| 1975 | EIF4B |
| 22827 | PUF60 |
| 7341 | SUMO1 |
| 1627 | DBN1 |
| 6629 | SNRPB2 |

| | |
|---|---|
| 54187 | NANS |
| 10480 | EIF3M |
| 7514 | XPO1 |
| 5496 | PPM1G |
| 4267 | CD99 |
| 7536 | SF1 |
| 813 | CALU |
| 27257 | LSM1 |
| 728689 | EIF3CL |
| 9520 | NPEPPS |
| 1347 | COX7A2 |
| 6566 | SLC16A1 |
| 8539 | API5 |
| 2271 | FH |
| 4676 | NAP1L4 |
| 3189 | HNRNPH3 |
| 3151 | HMGN2 |
| 4048 | LTA4H |
| 1977 | EIF4E |
| 5906 | RAP1A |
| 11052 | CPSF6 |
| 5049 | PAFAH1B2 |
| 5634 | PRPS2 |
| 56005 | MYDGF |
| 10465 | PPIH |
| 5162 | PDHB |
| 654483 | BOLA2B |
| 3033 | HADH |
| 6431 | SRSF6 |
| 11333 | PDAP1 |
| 23451 | SF3B1 |
| 2130 | EWSR1 |
| 29927 | SEC61A1 |
| 8664 | EIF3D |
| 6430 | SRSF5 |
| 128 | ADH5 |
| 1891 | ECH1 |
| 7323 | UBE2D3 |
| 84324 | SARNP |
| 79077 | DCTPP1 |
| 8661 | EIF3A |
| 85445 | CNTNAP4 |
| 5832 | ALDH18A1 |
| 5954 | RCN1 |

| | |
|---|---|
| 5908 | RAP1B |
| 4627 | MYH9 |
| 10959 | TMED2 |
| 11051 | NUDT21 |
| 4082 | MARCKS |
| 9948 | WDR1 |
| 57142 | RTN4 |
| 5917 | RARS |
| 3276 | PRMT1 |
| 29789 | OLA1 |
| 144983 | HNRNPA1L2 |
| 7458 | EIF4H |
| 55856 | ACOT13 |
| 8175 | SF3A2 |
| 85439 | STON2 |
| 9124 | PDLIM1 |
| 79084 | WDR77 |
| 6923 | TCEB2 |
| 38 | ACAT1 |
| 9217 | VAPB |
| 51637 | C14orf166 |
| 3646 | EIF3E |
| 27335 | EIF3K |
| 55832 | CAND1 |
| 10921 | RNPS1 |
| 10992 | SF3B2 |
| 317772 | HIST2H2AB |
| 80273 | GRPEL1 |
| 309 | ANXA6 |
| 55761 | TTC17 |
| 1892 | ECHS1 |
| 51386 | EIF3L |
| 5518 | PPP2R1A |
| 51128 | SAR1B |
| 56681 | SAR1A |
| 5716 | PSMD10 |
| 9521 | EEF1E1 |
| 6185 | RPN2 |
| 9377 | COX5A |
| 22826 | DNAJC8 |
| 8833 | GMPS |
| 7533 | YWHAH |
| 488 | ATP2A2 |
| 3735 | KARS |

| | |
|---|---|
| 1327 | COX4I1 |
| 81876 | RAB1B |
| 3704 | ITPA |
| 7165 | TPD52L2 |
| 3068 | HDGF |
| 6118 | RPA2 |
| 1431 | CS |
| 4735 | 45902 |
| 8761 | PABPC4 |
| 26973 | CHORDC1 |
| 10342 | TFG |
| 8662 | EIF3B |
| 16 | AARS |
| 58477 | SRPRB |
| 81 | ACTN4 |
| 5202 | PFDN2 |
| 10327 | AKR1A1 |
| 1350 | COX7C |
| 10627 | MYL12A |
| 84817 | TXNDC17 |
| 7905 | REEP5 |
| 5217 | PFN2 |
| 4698 | NDUFA5 |
| 5303 | PIN4 |
| 4942 | OAT |
| 387 | RHOA |
| 55611 | OTUB1 |
| 8667 | EIF3H |
| 2237 | FEN1 |
| 103910 | MYL12B |
| 6434 | TRA2B |
| 1356 | CP |
| 7385 | UQCRC2 |
| 2171 | FABP5 |
| 539 | ATP5O |
| 6117 | RPA1 |
| 6631 | SNRPC |
| 10952 | SEC61B |
| 30968 | STOML2 |
| 81542 | TMX1 |
| 7114 | TMSB4X |
| 3094 | HINT1 |
| 832 | CAPZB |
| 6723 | SRM |

| | |
|---|---|
| 6727 | SRP14 |
| 8880 | FUBP1 |
| 9230 | RAB11B |
| 5878 | RAB5C |
| 54732 | TMED9 |
| 11171 | STRAP |
| 23480 | SEC61G |
| 51493 | RTCB |
| 2108 | ETFA |
| 3150 | HMGN1 |
| 10204 | NUTF2 |
| 6182 | MRPL12 |
| 10598 | AHSA1 |
| 6426 | SRSF1 |
| 10487 | CAP1 |
| 8766 | RAB11A |
| 10236 | HNRNPR |
| 871 | SERPINH1 |
| 10063 | COX17 |
| 10539 | GLRX3 |
| 8666 | EIF3G |
| 136319 | MTPN |
| 7336 | UBE2V2 |
| 10146 | G3BP1 |
| 2224 | FDPS |
| 6282 | S100A11 |
| 8665 | EIF3F |
| 10923 | SUB1 |
| 57819 | LSM2 |
| 513 | ATP5D |
| 29097 | CNIH4 |
| 27258 | LSM3 |
| 1434 | CSE1L |
| 3185 | HNRNPF |
| 3028 | HSD17B10 |
| 5589 | PRKCSH |
| 29968 | PSAT1 |
| 1192 | CLIC1 |
| 471 | ATIC |
| 3149 | HMGB3 |
| 2950 | GSTP1 |
| 293 | SLC25A6 |
| 1329 | COX5B |
| 11335 | CBX3 |

| | |
|---|---|
| 6472 | SHMT2 |
| 6301 | SARS |
| 3692 | EIF6 |
| 7178 | TPT1 |
| 1994 | ELAVL1 |
| 231 | AKR1B1 |
| 79073 | TMEM109 |
| 59 | ACTA2 |
| 328 | APEX1 |
| 6625 | SNRNP70 |
| 25824 | PRDX5 |
| 873 | CBR1 |
| 4678 | NASP |
| 2109 | ETFB |
| 829 | CAPZA1 |
| 1854 | DUT |
| 6227 | RPS21 |
| 6633 | SNRPD2 |
| 72 | ACTG2 |
| 7086 | TKT |
| 8565 | YARS |
| 6627 | SNRPA1 |
| 11075 | STMN2 |
| 10728 | PTGES3 |
| 26517 | TIMM13 |
| 6888 | TALDO1 |
| 11315 | PARK7 |
| 9939 | RBM8A |
| 2098 | ESD |
| 23193 | GANAB |
| 7345 | UCHL1 |
| 10492 | SYNCRIP |
| 5094 | PCBP2 |
| 689 | BTF3 |
| 58527 | ABRACL |
| 5358 | PLS3 |
| 3148 | HMGB2 |
| 381 | ARF5 |
| 6741 | SSB |
| 9141 | PDCD5 |
| 10971 | YWHAQ |
| 10857 | PGRMC1 |
| 7529 | YWHAB |
| 3615 | IMPDH2 |

| | |
|---|---|
| 7919 | DDX39B |
| 7334 | UBE2N |
| 81611 | ANP32E |
| 375 | ARF1 |
| 2287 | FKBP3 |
| 5111 | PCNA |
| 1727 | CYB5R3 |
| 377 | ARF3 |
| 9446 | GSTO1 |
| 51155 | HN1 |
| 2280 | FKBP1A |
| 5464 | PPA1 |
| 6156 | RPL30 |
| 308 | ANXA5 |
| 1622 | DBI |
| 11021 | RAB35 |
| 3184 | HNRNPD |
| 10541 | ANP32B |
| 1917 | EEF1A2 |
| 8125 | ANP32A |
| 3012 | HIST1H2AE |
| 3013 | HIST1H2AD |
| 55766 | H2AFJ |
| 8329 | HIST1H2AI |
| 8330 | HIST1H2AK |
| 8331 | HIST1H2AJ |
| 8332 | HIST1H2AL |
| 8334 | HIST1H2AC |
| 8335 | HIST1H2AB |
| 8336 | HIST1H2AM |
| 85235 | HIST1H2AH |
| 8969 | HIST1H2AG |
| 92815 | HIST3H2A |
| 6432 | SRSF7 |
| 5037 | PEBP1 |
| 6647 | SOD1 |
| 3925 | STMN1 |
| 10209 | EIF1 |
| 811 | CALR |
| 7280 | TUBB2A |
| 9168 | TMSB10 |
| 347733 | TUBB2B |
| 345651 | ACTBL2 |
| 6234 | RPS28 |

| | |
|---|---|
| 9748 | SLK |
| 723790 | HIST2H2AA4 |
| 8337 | HIST2H2AA3 |
| 8338 | HIST2H2AC |
| 92521 | SPECC1 |
